# ST-GEARS: Advancing 3D Downstream Research through Accurate Spatial Information Recovery

**DOI:** 10.1101/2023.12.09.570320

**Authors:** Tianyi Xia, Luni Hu, Lulu Zuo, Yunjia Zhang, Mengyang Xu, Qin Lu, Lei Zhang, Lei Cao, Taotao Pan, Bohan Zhang, Bowen Ma, Chuan Chen, Junfu Guo, Chang Shi, Mei Li, Chao Liu, Yuxiang Li, Yong Zhang, Shuangsang Fang

**Affiliations:** BGI Research, Beijing 102601, China; BGI Research, Shenzhen 518083, China; BGI, Shenzhen 518083, China; BGI Research, Qingdao 266555, China; BGI Research, Wuhan 430074, China; Guangdong Bigdata Engineering Technology Research Center for Life Sciences, BGI research, Shenzhen 518083, China

## Abstract

Three-dimensional Spatial Transcriptomics has revolutionized our understanding of tissue regionalization, organogenesis, and development. However, to reconstruct single sections back to their *in situ* three-dimensional morphology, existing approaches either only adopt gene expression information to guide reconstruction or overlook shape correction against experiment-induced section distortions. This leads to significant discrepancies between reconstruction results and the actual *in vivo* locations of cells, imposing unreliable spatial profiles to downstream analysis. To address these challenges, we propose ST-GEARS (Spatial Transcriptomics GEospatial profile recovery system through AnchoRS), which solves optimized ‘anchors’ between *in situ* closest spots utilizing expression and structural similarity across sections and recovers *in vivo* spatial information under the guidance of anchors. By employing innovative Distributive Constraints into the Optimization scheme, it retrieves anchors with higher precision compared to existing methods. Taking these anchors as reference points, ST-GEARS first rigidly aligns sections, then introduces and infers Elastic Fields to counteract distortions. ST-GEARS denoises the fields using context information by Gaussian Denoising. Utilizing the denoised fields, it eliminates distortions and eventually recovers original spatial profile through innovative and mathematically proved Bi-sectional Fields Application. Studying ST-GEARS on both bi-sectional registration and complete tissue reconstruction across sectional distances and sequencing platforms, we observed its outstanding performance in spatial information recovery across tissue, cell, and gene levels compared to current approaches. Through this recovery, ST-GEARS provides precise and well-explainable ‘gears’ between *in vivo* situations and 3D *in vitro* analysis, powerfully fueling the potential of biological discoveries.

## Introduction

Spatial transcriptomics (ST) is an omics technology that fuels biological research based on measuring gene expression on each position-recorded spot across sliced tissues[1][2][3]. Notably, a range of methods has been developed. In situ sequencing (ISS)[4] platforms such as Barcoded Anatomy Resolved by Sequencing (BARseq)[5] and Spatially-resolved Transcript Amplicon Readout Mapping (STARmap)[6] rely on amplification, hybridization and imaging process to capture gene expression information. Next Generation Sequencing (NGS)[7] platform such as Visum[1], Stereo-seq[8] and Slide-Seq2[9] uses spatial barcoding and capturing in their implementations. These methods offer various sequencing resolutions ranging from 100 µm[10] to 500 nm[8], and can measure thousands[5] to tens of thousands[8] of genes simultaneously.

Single-slice ST studies have unleashed discoveries, and facilitated our understanding in diverse biological and medical fields[9][12][13][14][15]. Consequently, numerous processing pipelines and analysis models have been developed for ST data on a single section[16][17][18][19][20][21]. However, to truly capture transcriptomics in the real-world context, three-dimensional (3D) ST was designed to recover biological states and processes in real-world dimensions, without restriction of the isolated planes in single sectional ST studies. Various research has utilized the power of 3D ST to uncover insights in homeostasis, development, and diseases. Among them, Wang et al.[22] uncovered spatial cell state dynamics of Drosophila larval testis and revealed potential regulons of transcription factors. Mohenska et al.[23] revealed complex spatial patterns in murine heart and identified novel markers for cardiac subsections. And Vickovic et al.[24] explored cell type localizations in human rheumatoid arthritis synovium. The vast and large variety of downstream 3D research has posted the need for a reliable and automatic recovery method of *in vivo* spatial profile.

However, the collection process of ST data casts significant challenges onto the accurate reconstruction of 3D ST and the situation has not been overcome by current explorations. Specifically, in 3D ST experiments, individual slices are cross sectioned in a consistent direction, then manually placed on different chips or slides[14][25]. This operation introduces varying geospatial reference systems of distinct sections, and coordinates are distorted compared to their *in situ* states. The distortions occur due to squeezing and stretching effects during the picking, holding, and relocation of the sections. Such different geospatial systems and distortions complicates the recovery of *in situ* 3D profile. Among current recovery approaches, STUtility[26] realizes multi-section alignment through the registration of histology images, without considering either geospatial or molecular profile of mRNA, which leads to compromised accuracies. Recently published method PASTE[27], and its second version PASTE2[28] achieve alignment using both gene expression and coordinate information, through optimization of mapping between individual spots across sections. These methods cause inaccurate mappings and produces rotational misalignments due to the nonadaptive regularization factors, and their uniform sum of probability assigned to all spots upon presence of spots without actual anchors. All above approaches only consider rigid alignment, yet neglect the correction of shape distortions, resulting in shape inconsistency across registered sections. Published method Gaussian Process Spatial Alignment (GPSA)[29] considers shape distortions in its alignment, yet it doesn’t involve structural consistency in its loss function, which can cause the model to overfit to local gene expression similarities, leading to mistaken distortions of spatial information. Moreover, its hypothesis space involves readout prediction in addition to coordinates alignment, causing uncertainty in direction of gradient descent, and vulnerabilities to input noises. Another alignment approach, Spatial-linked alignment tool (SLAT)[30] also focuses on anchors construction between sections, yet it doesn’t provide a methodology to construct 3D transcriptomics profile. Other tools focus on analysis and visualization of 3d data, such as Spateo[31], VT3D[32] and StereoPy[33].

To address these limitations, we introduce ST-GEARS, a 3D geospatial profile recovery approach designed for ST experiments. By formulating the problem using the framework of Fused Gromov-Wasserstein (FGW) Optimal Transport (OT)[34], ST-GEARS incorporates both gene expression and structural similarity into the Optimization process to retrieve cross-sectional mappings of spots with the same *in situ* planar positions, also referred to as ‘anchors’. During this process, we introduce innovative Distributive Constraints that allow for different emphasis on distinct spot groups. The strategy addresses importance of expression consistent groups and suppresses inconsistent groups from imposing disturbances to optimization. Hence it increases anchor accuracy compared to current approaches. ST-GEARS utilizes the retrieved anchors to initially perform rigid alignment of sections. Subsequently, it introduces Elastic Field guided by the anchors to represent the deformation and knowledge to correct it according to each spot’s location. To enhance the quality of the field, Gaussian Smoothing is applied for denoising purposes. ST-GEARS then applies Bi-sectional Application to correction of each section’s spatial profile based on its denoised fields calculated with its neighboring sections. With validity proved mathematically, Bi-sectional Application eliminates distortions of sections, resulting in the successful recovery of a 3D *in situ* spatial profile.

To understand effects of ST-GEARS, we first studied its counterparts with innovations including anchors retrieval and elastic registration, respectively on human dorsolateral prefrontal cortex (DLPFC)[35], and Drosophila larva[22]. We found an advanced anchors accuracy of ST-GEARS compared to other available methods involving anchor’s concept and unveiled Distributive Constraints as reason behind the advancement. We validated the effectiveness of elastic registration process of ST-GEARS on both tissue shape smoothness and cross-sectional consistency. Then, we studied output of ST-GEARS and other methods on their reconstruction of Mouse hippocampus tissues[36], Drosophila embryo individual[22] and a complete Mouse brain[37]. The result was studied on morphological, cell and gene levels. ST-GEARS was found to be the only method that correctly reconstruct on all cases despite of cross-sectioning distance, number of sections, and sequencing platforms, and it was found to output the most accurate spatial information under both cell type labelling and hybridization evidence.

## Results

### ST-GEARS algorithm

ST-GEARS uses ST data as its inputs, including mRNA expression, spatial coordinates as well as approximate grouping information such as rough clustering and annotation of each observation. Then it recovers 3D geospatial profile in following steps (Fig. 1).

**Figure 1:**
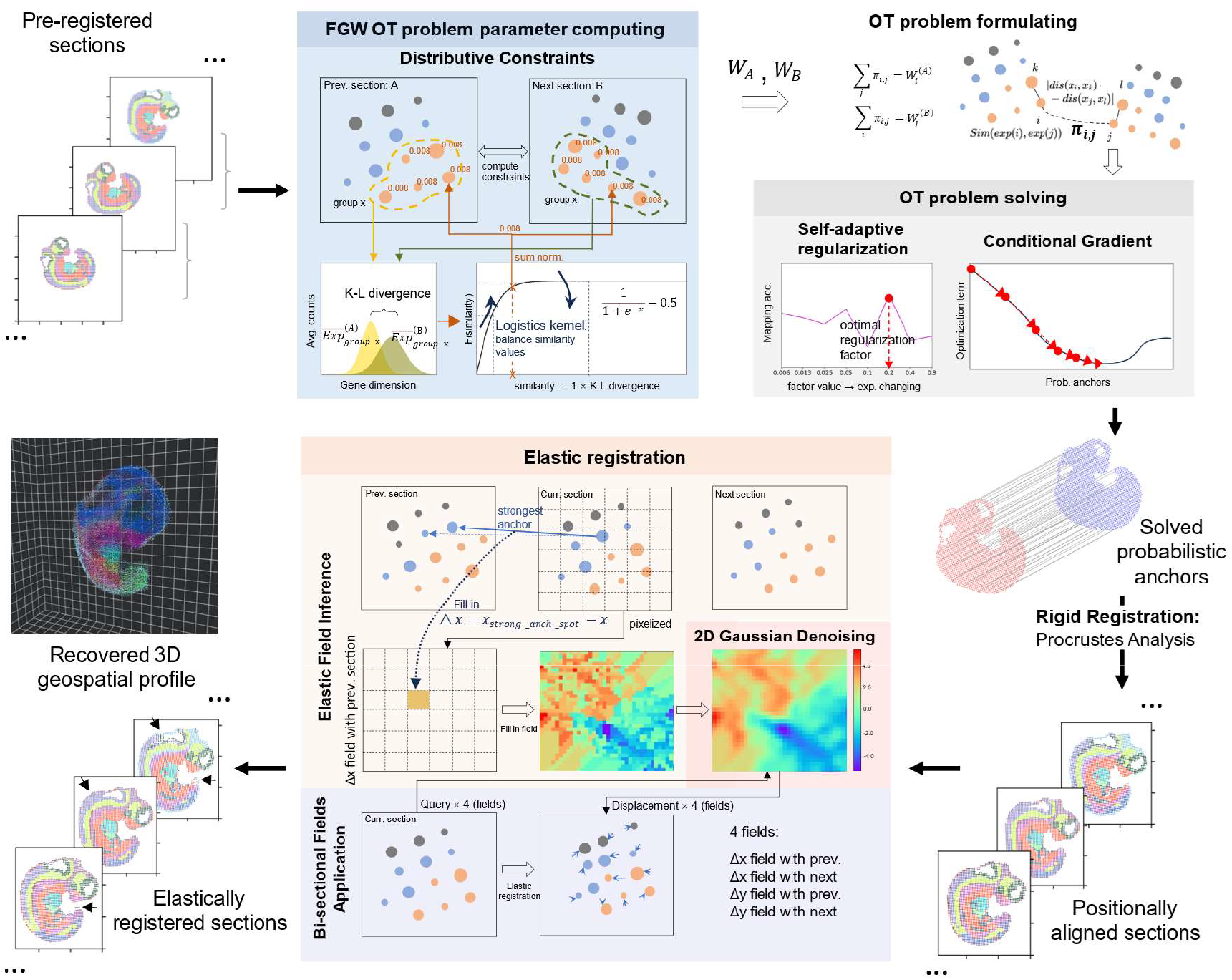
Three-Dimensional (3D) Spatial Transcriptomics (ST) Geospatial profile recovery with ST-GEARS. ST-GEARS recovers 3D in vivo spatial information through an automatic pipeline consisted by Fused Gromov Wasserstein (FGW) Optimal Transport (OT) problem formulating, OT problem solving, Procrustes Analysis, and distortion field computing & denoising. The input of the method is Unique molecular identifier (UMI) counts and location of each spot measured by ST technology, along with their annotations or cross-section clustering result. FGW OT finds probabilistic anchors joining spots with highest *in situ* proximity, through optimizing the combination of gene expression and structural similarity; the combined similarity visualization is adopted from Titouan et al.[34]. The OT problem is solved through Conditional Gradient (CG) method, leading to findings of the probabilistic anchors. Procrustes Analysis utilizes the anchors to solve optimal positional alignment between sections. Then the anchors information is utilized again to compute and denoise distortion field generated on each section, leading to the recovered 3D in vivo spatial information of the experimented tissue, or sample.

1. Optimization problem formulation under scheme of FGW OT. The formulation is established to enable solving of ‘anchors’, which are the joining of pair of spots with same *in situ* planar positions . Noticeably, each solved anchor is equipped with a probability that describes its strength of connection, and each spot is solved to have zero to multiple anchors. Among each two sections, section-specific groups of spots, and genes are initially excluded from the formulation to avoid their disturbances to anchors computing. Considering that connected spots are more spatially approximate, and more similar in gene expression because of shared cell identity[38][39], FGW was adopted to combine the gene expression and structural terms in optimization, which compensates inaccuracy and inadequacy of either information. Moreover, an innovative Distributive Constraints setting is designed and integrated into FGW OT’s formulation to relatively suppress the disturbances of groups of spots with lower cross-sectional expression similarity and to strengthen the influence of spots with higher similarity, leading to enhanced accuracy of anchor determination .
2. Optimization problem solving. Our designed Self-adaptive Regularization strategy automatically determines the weights of expression and structural terms in the optimization problem. This strategy leads to an optimal regularization factor across different section distances, spot sizes, extent of distortions, and data quality such as level of diffusion. Conditional Gradient[34] is adopted as optimizer, which updates anchors iteratively towards higher expression and structural similarity with each iteration. The efficacy of Conditional Gradient has been demonstrated through its convergence to a local optimal point[40], thereby ensuring the robustness and effectiveness of our approach.
3. Rigid registration by Procrustes Analysis[41]. After filtering out anchors with relatively low probabilities, the optimal transformation and rotation of each section are analytically solved through Procrustes Analysis, which minimizes summed spatial distances of spots anchored to each other. With the transformation and rotation applied, sections are positionally aligned.
4. Elastic registration guided by anchors. On each rigidly registered section, our innovated Elastic Field Inference leverages the anchored spots on its front and next sections as guidance to correct deformation. It transforms offsets between spots into deformation fields between sections, which meanwhile enables data quality enhancement by 2D Gaussian Denoising. Making use of continuity of deformation at local scales, 2D Gaussian Denoising convolutes all over the fields to reduce noises caused by any inaccurately generated anchors. With denoised fields, our designed Bi-sectional Fields Application corrects each section’s deformation according to its fields calculated with front and next sections. The bi-sectional correction method is mathematically proved to approximately recover each section’s spatial profile to its original state.

### Enhancement of anchor retrieval accuracy through Distributive Constraints

As was unfolded, ST-GEARS is an algorithm flow jointly constituted of probabilistic anchor computation and spatial information recovery. Hence, to validate the effectiveness of our method and demonstrate its underlying design philosophy, we conducted comprehensive studies on the two counterparts using real-world data. To begin, we utilized the DLPFC dataset[35] to study our anchors retrieving accuracy with emphasis on the effect of Distributive Constraints design.

To assess the effects of Distributive Constraints on anchor accuracy, we compared ST-GEARS with and without this setting, and with other constraints involving methods including PASTE and PASTE2. We investigated constraint values assigned by these methods, as well as their solved number of anchors and maximum anchor probability of each spot. Furthermore, we examined the cell types that were considered connected based on the computed anchors to assess accuracy of anchors. Among the methods we compared, ST-GEARS with Distributive Constraints was found to assign different constraint values to spots within different neuron layers, while the others assigned uniform constraints to all layers (Fig. 2a, Supplementary Fig. 1). The results of ST-GEARS showed that both number of anchors and the anchors’ maximum probabilities for each spot were lower in Layer 2 and Layer 4 compared to the thicker layers. However, this pattern was not observed in methods without Distributive Constraints setting (Fig. 2a, Supplementary Fig. 1). To illustrate the impact of this strategy on anchor accuracy, we tagged each spot with annotation of its connected spot by anchor with highest probability. We then compared this result to the tagged spot’s original annotation (Fig. 2a, Supplementary Fig. 1). Under Distributive Constraints, ST-GEARS achieved a significantly higher proximity between annotations compared to PASTE and our method without Distributive Constraints. PASTE2 also led to approximate annotations, but it anchored multiple spots to spots from neighboring layers, particularly those near layer boundaries. To evaluate the precision of anchors, we conducted a comparison with the Mapping accuracy index introduced by PASTE[27]. This index measures the weighted percentage 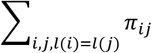 of anchors that connect spots with same annotation. As a result, ST-GEARS outperformed PASTE2, and reached a score that was over 0.5 (out of 1) higher than both PASTE and our method without Distributive Constraints (Fig. 2a, Fig. 2b, Supplementary Fig. 1).

**Figure 2:**
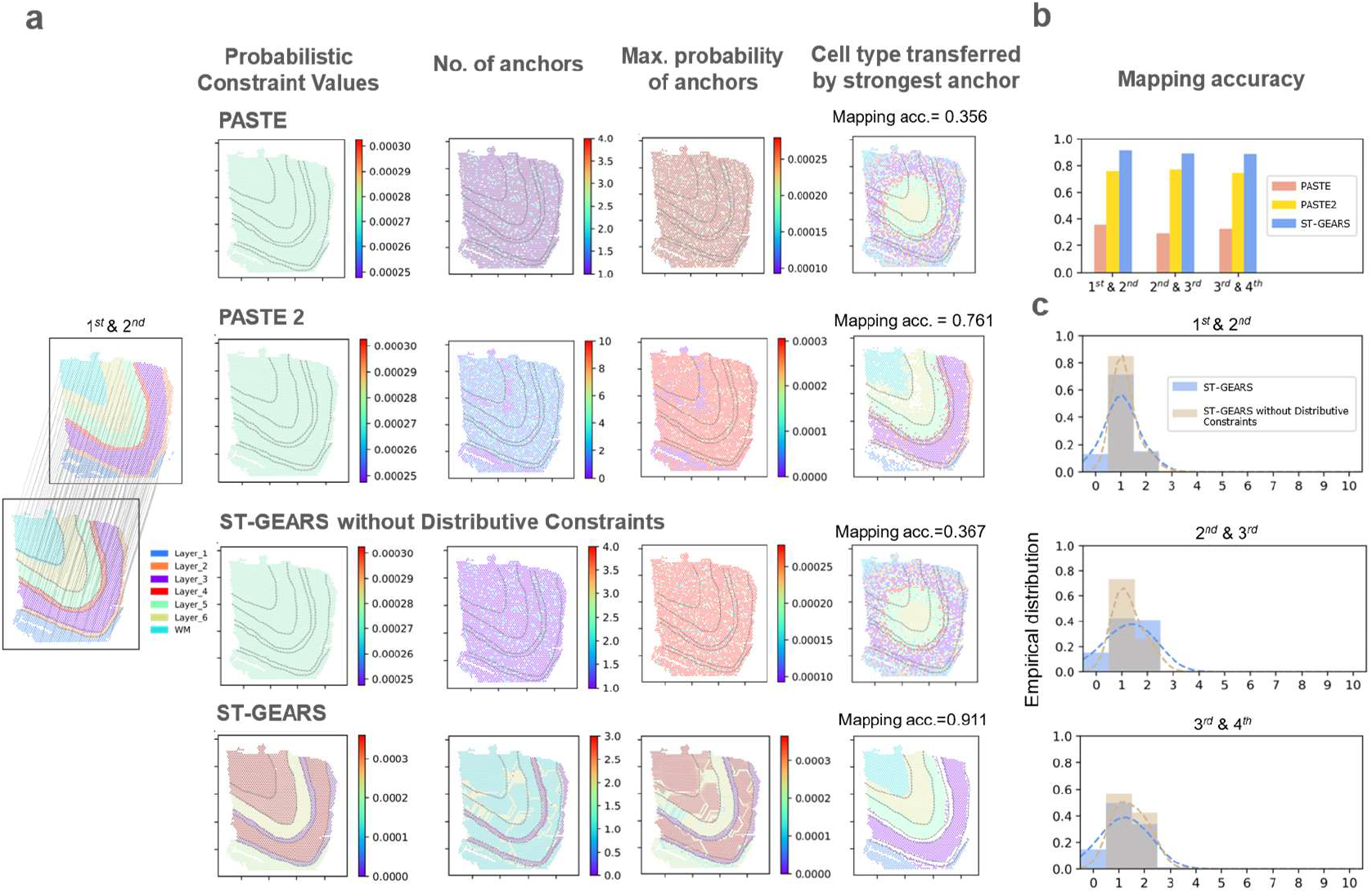
Demonstration of anchors accuracy by ST-GEARS. (a) (from left to right) 1^st^ and 2^nd^ human dorsolateral prefrontal cortex (DLPFC) section of patient #3 by Maynard et al.[35] with their provided annotations and our anchors showcase, (of the same section pair) probabilistic constraints settings in Optimal Transport (OT) problem formulating, no. of anchors computed on each spot, max. anchor probability value computed of each spot, and annotated cell type mapped back to spots through computed anchors; (from top to bottom) respectively by PASTE, PASTE2, ours without distributive constraints setting, and ours. The distinction of different cell types on the 1st section is marked by dotted lines. Mapping accuracy is marked alongside respective cell type mapping visualizations. (b) Mapping accuracy measured on anchors of sections pairs used in (b) by PASTE, PASTE2 and ST-GEARS. (c) Comparison of no. of anchors histograms between ST-GEARS and ST-GEARS without distributive constraints, of sections pairs of 1^st^ & 2^nd^, 2nd & 3^rd^, and 3^rd^ & 4^th^ sections. The Probability Density Function (PDF) estimated by Gaussian kernel was plotted in dotted lines with the same color of histograms, to highlight the distribution differences.

To uncover the reasons behind the aforementioned phenomena, as the functional area in between thicker neocortical layers, thinner neocortical layers have comparable transcriptomic similarity with their adjacent layers in gene expression, than with its own cell type[35][44]. This implies that, in contrast to thicker layers, thinner layers tend to introduce more disturbances during anchor computation. However, the Distributive Constraints imposed suppression on these cell types by assigning a smaller sum of probability to each of their spots. The suppression was reflected in above results where each spot in Layer 2 and Layer 4 has fewer assigned anchors and a lower maximum probability (Fig. 2a, Supplementary Fig. 1). Further analysis on all spots in the DLPFC reveals that a certain percentage of spots were suppressed in anchor generation due to the Distributive Constraints (Fig. 2c, Supplementary Fig. 2).

### Recovery of *in situ* shape profile through elastic registration

We then utilized Drosophila larva data to investigate the spatial profile recovery effect of ST-GEARS, with an emphasis on our innovated elastic registration. We first applied rigid registration to Drosophila larva sections and observed a visually aligned configuration of individual sections (Supplementary Fig. 3). By further mapping cell annotations back to their previous sections, according to the strongest anchors of each spot, the projected annotations are visually in match with original ones (Supplementary Fig. 4). The accuracy of the mapping matching between annotations was quantified by Mapping accuracy (Supplementary Fig. 5). The above findings validated that ST-GEARS produced reliable anchors and accurately aligned sections through rigid registration. However, when stacking the sections together, we observed an inconsistency on the edge of lateral cross-section of the rigid result (Supplementary Fig.6). This inconsistency doesn’t conform to the knowledge of intra-tissue and overall structural continuity of Drosophila larvae.

After applying elastic registration to the rigidly-aligned larva, we observed a notable improvement in the continuity of the cross section above, indicating a closer-to-real spatial information being retrieved. To further understand the effect of elastic operation on the dataset, we compared the changes in area of the complete body and three individual tissues (trachea, central nervous system (CNS), and fat body) on all sections. We observed an enhanced smoothness in the curves of elastically registered sections, which aligns with the continuous morphology of the larva as expected by theoretical knowledge . To quantify the smoothing effect, we calculated Scale-independent Standard Deviation of Differences (*SI-STD-DI* = *STD*({*s*_*i*_ − *s*_*i*-1_: *i* ∈ [1,2, …, *I* − 1]})/ |*mean*({*s*_*i*_ − *s*_*i*-1_: *i* ∈ [1,2, …, *I* − 1]})|) onto the curves, which measures the smoothness of area changes along the sectioning direction (Fig. 3a and Methods). A decrease of SI-STD-DI on all tissues and the body provided empirical evidence for the improved smoothness. To further investigate the recovery of internal structures, we introduced Mean Structural Similarity (MSSIM). MSSIM takes structurally consistent sections as input, and measures pairwise internal similarity of reconstructed result using annotations or clustering information (Supplementary Fig. 7). (See Methods for details.) An improved MSSIM was noticed on all 4 sections, indicating that elastic registration further recovers internal geospatial continuity on basis of rigid operation(Fig. 3b). By comparing registration effect of individual sections, we also observed that the elastic process successfully rectified a bending flaw along the edge of the third section, (Fig. 3c). The shape fixing highlighted that ST-GEARS not only yielded a more structurally consistent 3D volume, but also provided a more accurate morphology for single sections. The improved smoothness, the recovered structural continuity, and the shape fixing collectively demonstrate that elastic registration effectively recovers geospatial profile.

**Figure 3:**
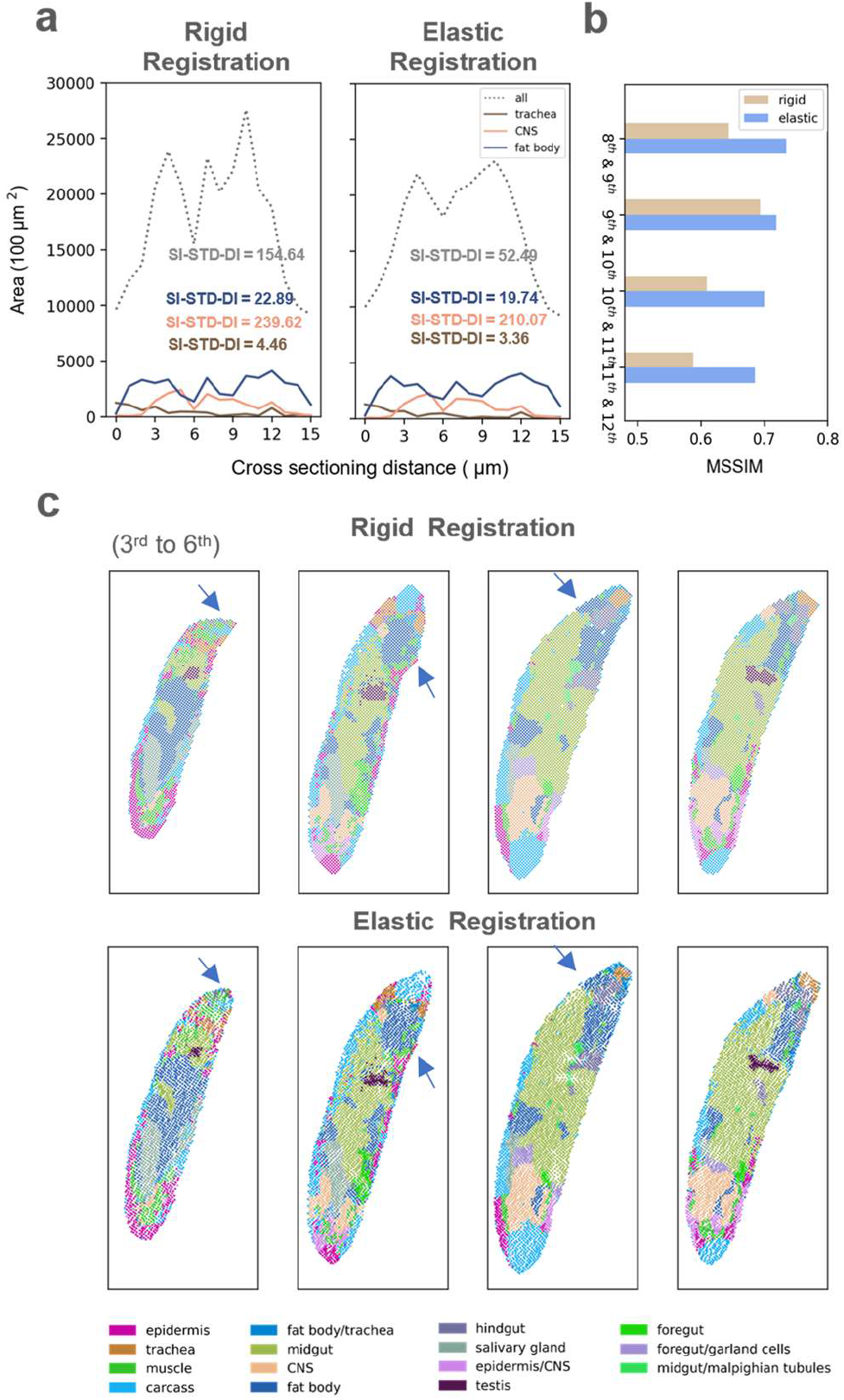
Demonstration of spatial information recovery effect of elastic registration of ST-GEARS. (a) A comparison of area changes of 3 tissues and complete body of Drosophila Larva, between result of rigid registration and result of elastic registration appended to rigid registration. The areas are calculated based on recovered spot position of different tissues along cross-sectioning direction. Standard Deviation of Differences (SI-STD-DI) quantifying the smoothness is marked alongside each curve. (b) A comparison of Mean Structural Similarity (MSSIM), of selected section pairs from Drosophila Larva (L3), between result of rigid registration only and result of elastic registration appended to rigid registration. The chosen section pairs are the structurally consistent ones. (c) Comparison of individual sections recovered by rigid registration only and by elastic registration appended to rigid registration, of 1^st^ to 5^th^ section of Drosophila Larva (L3). Shape correction of bended area in the 3^rd^ section, and increased cross-sectional consistency on the 4^th^ and 5^th^ section were highlighted by blue arrows.

With elastic process validated and applied onto rigid registration result, the recovery of spatial information was completed. Stacking individual sections of the elastic result, a complete geospatial profile of the larva was generated (Supplementary fig. 8), visualizing the ST-GEARS’ ability of *in situ* spatial information recovery.

### Application to sagittal sections of Mouse hippocampus

After validating the component phases of ST-GEARS, we proceeded to apply the method to multiple real-world problems to recover geospatial profiles. We first focused on two sagittal sections of mouse hippocampus[36] (Supplementary Fig. 9) that were 10 μm apart, accounting for 1-2 layers of Cornu Ammonis (CA) 1 neurons[45]. Considering the proximity of these sections, we assumed no structural differences between them.

To compare the differences of registration effect among methods, we extracted CA fields and dentate gyrus (DG) beads (Supplementary Fig. 10), then stacked the two sections for a more obvious contrast (Fig. 4b). PASTE2 failed in performing the registration, leaving the sections unaligned. By GPSA, the sections’ positions were aligned, yet the 2^nd^ section were squeezed into a narrower region than first one, leading to a contradiction of region’s location. The ‘narrowing’ phenomena may be caused by the overfitting of GPSA model on gene expression similarity, since it doesn’t involve structural similarity between registered sections in loss function. The scale on horizontal and vertical axis was distorted due to the equal scale range strategy adopted in GPSA’s preprocessing. In the comparison between PASTE and ST-GEARS, our method demonstrates a more accurate centerline overlapping of CA fields and DG compared to PASTE. This indicated an enhanced recovery of spatial structure consistency and an improved registration effect. To quantitatively evaluate these findings, we utilized the MSSIM index as a measure of structural consistency and compared it among PASTE, PASTE2, GPSA and ST-GEARS (Fig. 4a). Consistent with the results of centerline, ST-GEARS achieved a higher MSSIM score than GPSA and PASTE, surpassing PASTE2 by more than 0.2 out of 1.

**Figure 4:**
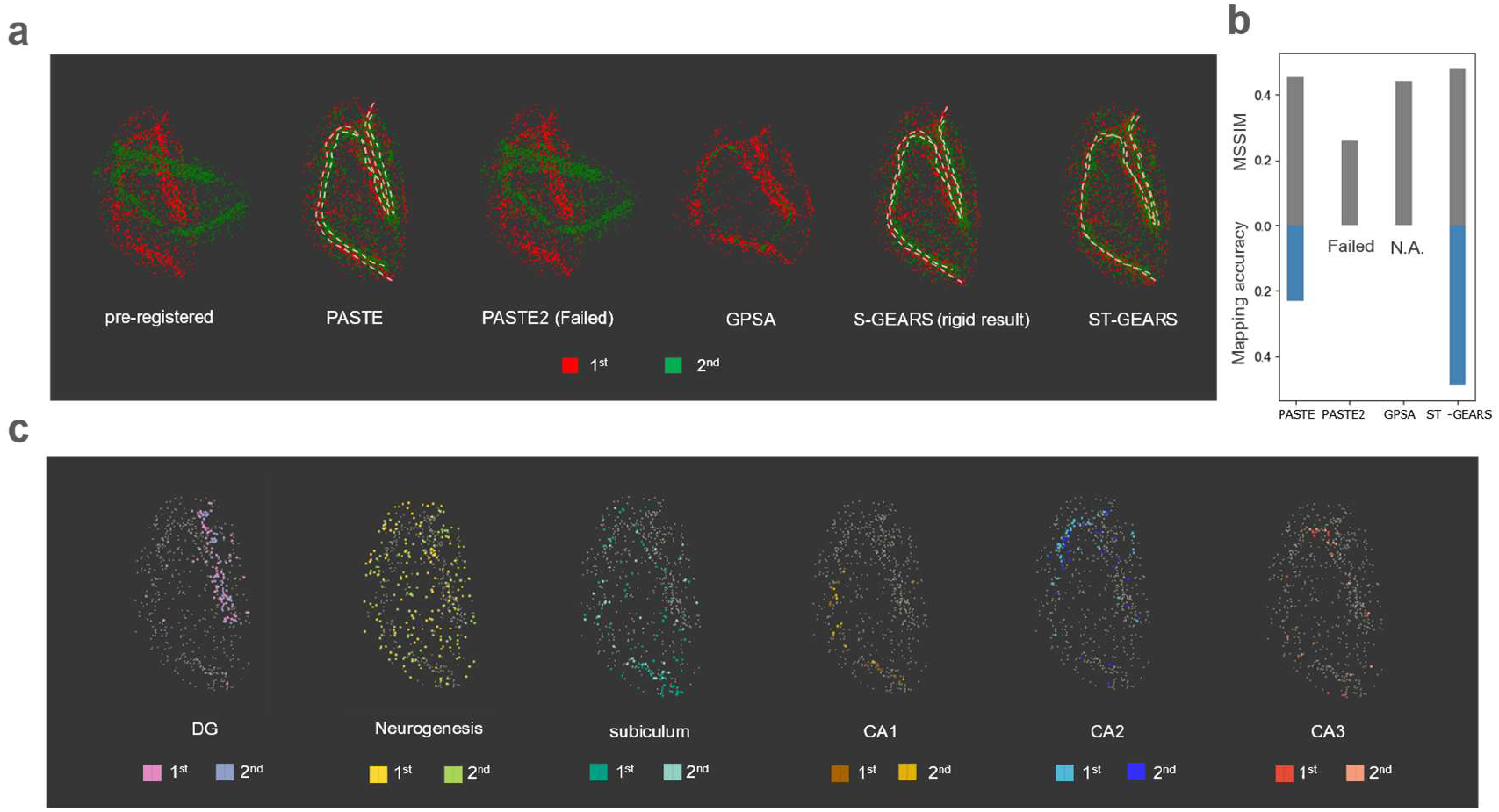
Registration of Mouse hippocampus, respectively by PASTE, PASTE2, GPSA and ST-GEARS. (a) Stacked projections of Cornu Ammonis (CA) fields and dentate gyrus (DG), of pre-registered and registered result of Mouse hippocampus sagittal sections with 10 µm distance, respectively by PASTE, PASTE2, GPSA and ST-GEARS. (b) A comparison of both MSSIM and Mapping accuracy of the 2 registered sections, across PASTE, PASTE2, GPSA and ST-GEARS. (c) Stacked projections of region-specific cell types including DG, Neurogenesis, subiculum, CA1, CA2 and CA3, registered by ST-GEARS. Each column highlights the stacked projection of a single cell type.

To understand reasons behind our enhancement, we thoroughly examined the anchors generated by PASTE, PASTE2 and ST-GEARS, as well as the effects of our elastic registration. By mapping cluster information of the 2^nd^ section to the 1^st^, and the 1^st^ to the 2^nd^ through anchors, we found correspondences between the projected and original annotations (Supplementary Fig. 11). Accordingly, our Mapping accuracy was over 0.25 higher than PASTE and over 0.45 than PATSE2 (Fig. 4a), indicating our exceptional anchor accuracy. To understand and further substantiate this advantage, we visualized the probabilistic constraints and its resulted anchors probabilities (Supplementary Fig. 12a). It is worth noting that ST-GEARS implemented cell-type-specific constraints, in contrast to the uniform distributions used by PASTE. As a result, a certain percentage of spots were found to be suppressed in anchors connection by ST-GEARS (Supplementary Fig. 12b) compared to PASTE, leaving the registration to rely more on spots with higher cross-sectional similarity and less computational disturbances, and hence lead to a higher anchor accuracy. In the study of elastic effect, we found an increased overlapping of centerlines by elastic registration than by rigid operation only when overlapping CA fields and DG (Fig. 4b). Quantitively by MSSIM, the cross-sectional similarity was found to be increased by elastic registration (Supplementary Fig. 13). These findings suggest that the combination of Distributive Constraints and elastic process contributed to the enhanced registration of the Mouse hippocampus.

To explore the potential effect of impact of our registration on downstream analysis, we extracted region-specific cell types from the sections, and analyzed their overlapping through stacking registered sections together (Fig. 4c). In all cell types including DG, Neurogenesis, subiculum, CA1, CA2 and CA3, the distribution regions from both sections were nearly identical. The overlapping result unveils that ST-GEARS integrated the spatial profile of same cell subpopulations, enabling a convenient and accurate downstream analysis of multiple sections.

### Application to 3D reconstruction of Drosophila embryo

Besides tissue level registration of Mouse hippocampus, to evaluate the performance of ST-GEARS in reconstructing individual with multiple sections, we further tested it on a Drosophila embryo. The transcriptomics of embryo was measured by Stereo-seq, with 7 μm cross-sectioning distance[22]. By quantifying the registration effect of spatial information recovery and comparing it to PASTE, PASTE2 and GPSA, we found that ST-GEARS achieved the highest MSSIM in five out of the six structurally consistent pairs (Fig. 5a). On the pair where ST-GEARS did not result in highest MSSIM, it surpassed PASTE, and achieved a similar score to PASTE2. By comparing area changes with SI-STD-DI quantification of the complete section, and three individual tissues including epidermis, midgut and foregut, ST-GEARS yielded higher smoothness on all regions than all other approaches, both visually and quantitatively(Fig. 5b).

**Figure 5:**
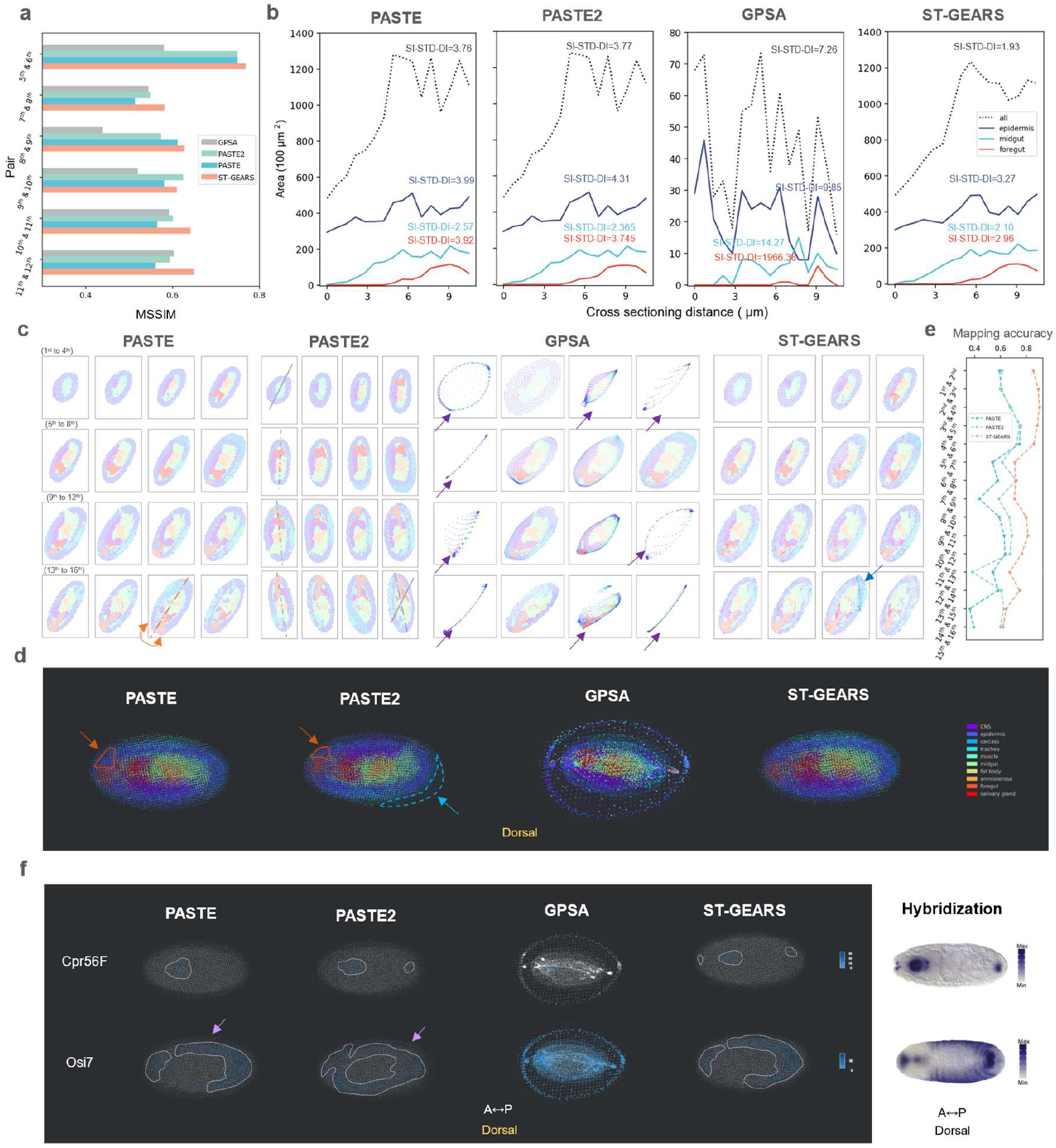
Three-Dimensional (3D) reconstruction of Drosophila Embryo, respectively by PASTE, PASTE2, GPSA and ST-GEARS. (a) A comparison of Mean Structural Similarity (MSSIM), of section pairs that are structurally consistent from Drosophila Embryo (E14-16h), between reconstruction result of PASTE, PASTE2, GPSA and ST-GEARS. (b) A comparison of area changes of 3 tissues and complete body of Drosophila Embryo, along cross-sectioning direction, between reconstruction result of PASTE, PASTE2, GPSA and ST-GEARS. Standard Deviation of Differences (SI-STD-DI) is marked alongside each curve to quantify the smoothness. (c) Reconstructed individual sections with recovered spatial location of each spot. In result of PASTE, the incorrect flipping on the 15^th^ section was highlighted in orange. In result of PASTE2, gradual rotations were marked by the 1^st^, 5^th^, 9^th^, 13^th^ and 16^th^ sections’ approximate symmetry axis whereas symmetry axis of the 1^st^ section was replicated onto the 16^th^ for angle comparison. In result of GPSA, mistakenly distorted sections were marked by purple arrows. In result of ST-GEARS, the fix of dissecting area on the 15^th^ section was marked by a blue arrow. (d) Dorsal view of 3D reconstructed Drosophila embryo by PASTE, PASTE2, GPSA and ST-GEARS. The inaccurate regionalization of midgut was circled and pointed with arrow in orange. The resulted extruding part of single section by PASTE2 was circled and pointed in blue. (e) Mapping accuracy of all section pairs by PASTE, PASTE2 and ST-GEARS. (f) By dorsal view, regionalization of marker gene Cpr56F and Osi7 by PASTE, PASTE2, GPSA and ST-GEARS, and their comparison with hybridization result from Berkeley Drosophila Genome Project (BDGP) database. The gathering expression regions were highlighted by dotted lines.

To compare the reconstruction effect, we studied both registered individual section, and reconstructed 3D volume. Among the methods compared, PASTE produced a wrong flipping on the 15^th^ section along A-P axis (Fig. 5c). Stacking sections back to 3D and investigating on dorsal view, the wrong flipping caused a false regionalization of foregut circled in orange (Fig. 5d). Along the first to last section registered by PASTE2, a gradual rotation was witnessed (Fig. 5c), leading to over 20 degrees of angular misalignment between the first and the last section. Similar to PASTE, this misalignment also caused the wrong regionalization of foregut in 3D map (Fig. 4d). Equally induced by the rotation, sections were found to extrude in the 3D result circled in blue, breaking the round overall morphology of the embryo. GPSA caused false distortion of 8 out of 16 sections as pointed by purple arrows (Fig. 5c) and the stacked sections formed a dorsal view of an isolated circle and an inner region (Fig. 5d). The phenomena may be due to its overfitting onto expressions, which is caused by the contradiction between its hypothesis of consistent readout across sections, and the large readout variation across 16 sections in this application. In contrast, ST-GEARS avoided all of these mistakes in its results (Fig. 5c). From the perspective of individual section profiles, noticeably in the 15th section, we observed a significant reduction in the dissecting region between two parallel lines, indicating the successful fixation of flaws in the session.

To comprehend the rationale behind our improvement, we analyzed the anchors generated by the three methods and the impact of our elastic registration. In the investigation of anchor accuracy, we discovered that ST-GEARS achieves the highest mapping accuracy among all section pairs (Fig. 5e), suggesting its advanced ability to generate precise anchors, which forms the basis for precise spatial profile recovery. To understand this advancement, probabilistic constraints and its resulted anchors distributions (Supplementary Fig. 14, Supplementary Fig. 15) were studied. With Distributive Constraints (Supplementary Fig. 14a), ST-GEARS generated different maximum probabilities on different cell types (Supplementary Fig. 14b), which indicates that cell types with higher cross-sectional consistency were prioritized in anchor generation. This selection led to reduced computational disturbances, and hence higher accuracy of anchors. In study of elastic registration in shape smoothness, we witnessed an increased level of smoothness of tissue epidermis, foregut, and midgut, as well as the complete section, through area changes quantified by SI-STD-DI index (Supplementary Fig. 16). In internal structure aspect, an increased MSSIM of structural consistent pairs were noticed (Supplementary Fig. 17). An experimental flaw on the 15^th^ section was also fixed by elastic registration (Supplementary Fig. 18). Above findings point that the enhancement of registration accuracy on Drosophila embryo was induced by Distributive Constraints and elastic process.

By mapping spots back to 3D space, we further investigated the effect of different method on downstream analysis, in the perspective of genes expression (Fig. 5f). Cpr56F and Osi7 were selected as marker genes, which were found to respectively highly express in foregut, and foregut plus epidermis region[22]. Investigating Cpr56F expression by ST-GEARS from dorsal view, we noticed three highly expressing regions, at anterior end, front region, and posterior end of the embryo. The finding matches the hybridization result of stage 13-16 Drosophila embryo extracted from Berkeley Drosophila Genome Project (BDGP) database. In contrast, none of PASTE, PASTE2 and GPSA presented high expression at all three locations. When analyzing the distribution of Osi7 by PASTE, we noticed a sharp decrease in expression from inner region to the outer layer, contradicting the prior knowledge of high expression in the epidermis. Similarly, PASTE2 failed to capture expression in outer layers and instead revealed a high expression in one inter-connected area, which did not correspond to the separate expression regions observed in hybridization result. No spatial pattern was witnessed when analyzing distribution of Osi7 by GPSA, which forms an obvious contrast to its hybridization evidence. Comparably, none of the violations was shown in the result of ST-GEARS. The comparison of spatial distribution indicated our potential capability to better enhance the process of downstream gene-related analysis.

- **Application to Mouse brain reconstruction** The design of 3D experiments involves various levels of sectioning distances[22][36][37]. To further investigate the applicability of ST-GEARS on ST data with larger slice intervals, we applied the method to a complete mouse brain hemisphere dataset, which consists of 40 coronal sections (Supplementary Fig. 19a), with a sectioning distance of 200 μm[37]. The transcriptomics data was measured by BARseq, which includes sequencing data and its cross-modal histology images. Each observation represents captured transcriptomics surrounded by the boundary of a cell.

Through respectively applying PASTE, PASTE2, GPSA and ST-GEARS onto the dataset, we observed multiple misaligned sections produced by approaches including PASTE, PASTE2 and GPSA (Supplementary Fig. 19b, Supplementary Fig. 19c, Supplementary Fig. 19d, Fig. 6a). In PASTE, these misalignments include 2 sections with approximately 180° angular misalignment (Supplementary Fig. 19b), . By PASTE2, 4 rotational misalignments and 8 positional misalignments were noticed (Supplementary Fig. 19d). By GPSA, 12 sections were observed to be rotationally misaligned, and 3 sections were mistakenly distorted (Supplementary Fig. 19b), probably due to its overfitting onto expressions discussed in analysis of Drosophila embryo. The scale on horizontal and vertical axis was distorted maybe due to the similar reason analyzed in Mouse hippocampus. As a clear contrast, our algorithm correctly aligned all 40 sections with 200 μm intervals (Supplementary Fig. 19e). To more accurately assess the result of our registration, we employed the direction of the cutting lines induced during tissue processing[37], and compared the consistency of tilt angles of these lines in the 20^th^, 25^th^, 26^th^, 27^th^, 33^rd^, 34^th^ and 37^th^ slices where these lines are visible. Notably, neither visual angle differences nor cutting line curving were observed, indicating that the sections were properly aligned by ST-GEARS (Fig. 6e, Supplementary Fig. 19d). To quantify the registration accuracy in aspect of structural continuity, we calculated MSSIM scores of 11 section pairs that are structural consistent (Fig. 6b). Consistent with the visual observations, PASTE2 presented a much larger score range than other methods, which reflects its instability across sections in this dataset, and GPSA exhibited the lowest median MSSIM score indicating its suboptimal average performance. By comparison, PASTE yielded a higher median score and a smaller variation, while ST-GEARS resulted in the highest median score and the smallest variation among all methods.

**Figure 6:**
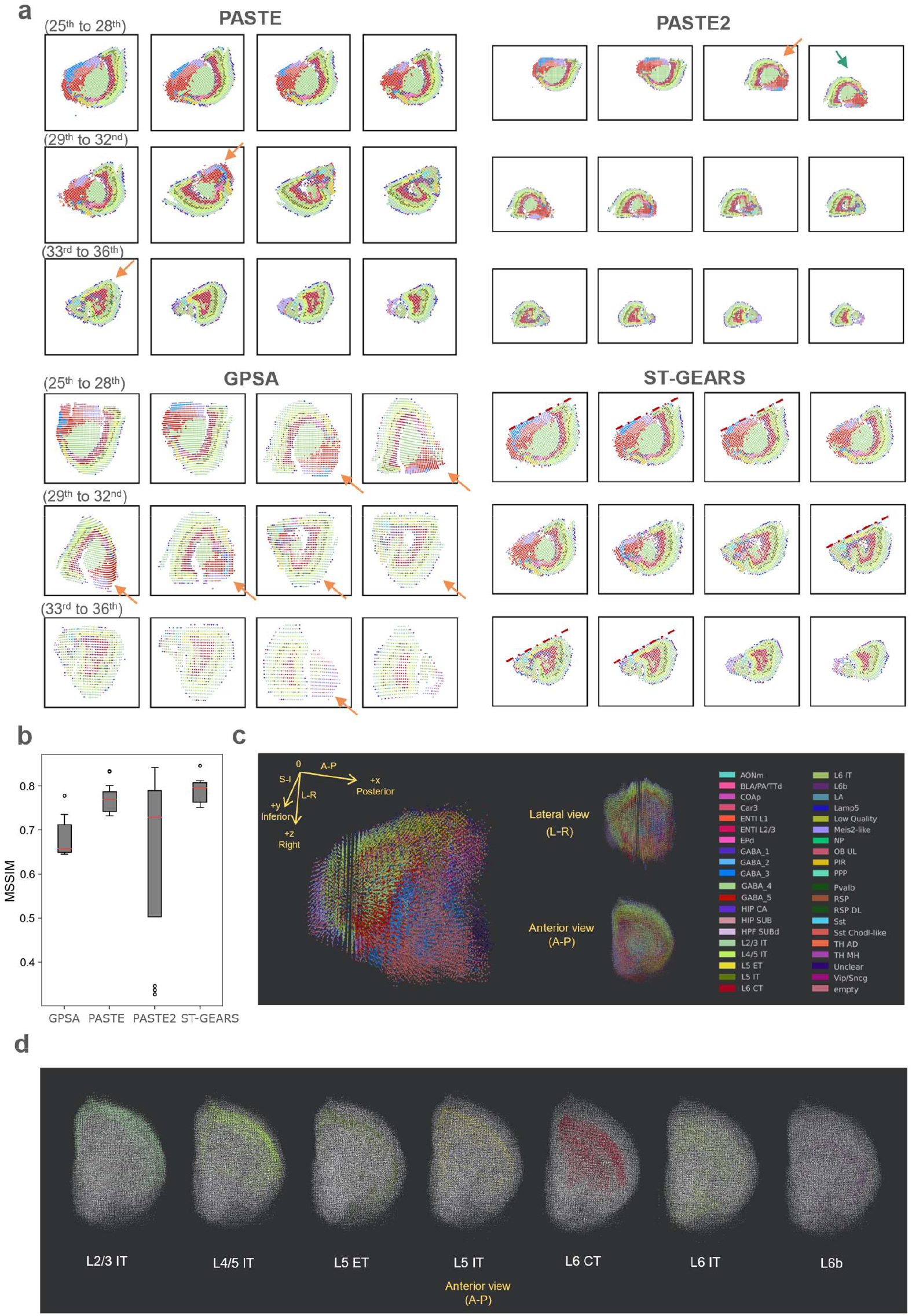
Three-Dimensional (3D) reconstruction of Mouse Brain, respectively by PASTE, PASTE2, GPSA and ST-GEARS. (a) Reconstructed individual sections with recovered spatial location of each spot from the 25^th^ to 36^th^ section. Positional misalignments are marked by arrows of green, and angular misalignments are marked by arrows of orange. Visible cutting lines by ST-GEARS are marked by dotted lines. (b) A comparison of Mean Structural Similarity (MSSIM) score of section pairs that are structurally consistent, between result of PASTE, PASTE2, GPSA and our method. The red lines positions show median score; the box extends from the first quartile (Q1) to the third quartile (Q3) of scores; the lower whisker is at the lowest datum above Q1 – 0.5*(Q3-Q1), and the upper whisker is at the highest datum below Q3 + 0.5*(Q3-Q1); scores out of whiskers range are marked by circles. (c) Perspective, Lateral and Anterior view of reconstructed mouse brain hemisphere. (d) Anterior view of layer cell types distribution of reconstructed mouse brain hemisphere.

To understand the reasons behind our progress, we examined anchor accuracy changes with regularization factors during ST-GEARS computation (Supplementary Fig. 20). Out of 39 section pairs, we observed a change in mapping accuracy greater than 0.1 (out of 1) in 12 pairs. By Self-adaptive Regularization which was designed to face varying data characteristics which also includes varying section distances, regularization factor that leads to optimal mapping accuracy was selected, leading to an increased anchors accuracy in the 12 section pairs. Notably, among these 12 pairs, pairs 29^th &^ 30^th^, 31^st^ & 32^nd^ and 32^nd^ & 33^rd^ were correctly aligned by ST-GEARS but misaligned by PASTE, which doesn’t adopt any self-adaptive regularization strategy.

After validating the registration result, we investigated the recovered cell-types’ distribution in the 3D space to assess the effectiveness of the reconstruction and its impact on further analysis. We observed that the complete morphology of hemisphere was recovered by ST-GEARS, with clear distinction of different tissues on perspective, lateral and anterior views (Fig. 6c). We further studied the distribution of separate cell types within cortex layers and found that 3D regionalization of each cell type was recovered by ST-GEARS (Fig. 6d). The reconstructed result indicated the adaptability of ST-GEARS across various scales of sectioning intervals, and its applicability on both bin-level, and cell-level datasets on which histology information is incorporated.

## Discussion

We introduce ST-GEARS, a 3D geospatial profile recovery approach for ST experiments. Leveraging the formulation of FGW OT, ST-GEARS utilizes both gene expression and structural similarities to retrieve cross-sectional mappings of spots with same *in situ* planar coordinates, referred to as ‘anchors’. To further enhance accuracy, it uses our innovated Distributive Constraints to enhance the accuracy. Then it rigidly aligns sections utilizing the anchors, before finally eliminating section distortions using Gaussian-denoised Elastic Fields and its Bi-sectional Application.

We validate counterpart of ST-GEARS including anchors retrieval and elastic registration, respectively on DLPFC and Drosophila larva dataset. In the validation of anchors retrieval, through Mapping accuracy evaluation of retrieved anchors, ST-GEARS consistently outperformed PASTE and PASTE2 across all section pairs. We show Distributive Constraints as reasons behind its distinguished performance, which effectively suppressed the generation of anchors between spot groups with low cross-sectional similarity while enhances their generation among groups with higher similarity. To investigate the effectiveness of the elastic registration process, we evaluate the effects of tissue area changes and cross-sectional similarity using the Drosophila larvae dataset. Both smoother tissue area curves and higher similarity observed between structurally consistent sections confirm the efficacy of the elastic process of ST-GEARS.

We demonstrate ST-GEARS’s advanced accuracy of reconstruction compared to current approaches including PASTE, PASTE2 and GPSA, and its positive impact on downstream analysis compared to existing approaches. Our evaluation encompasses diverse application cases, including registration of two adjacent sections of Mouse hippocampus tissue measured by Slide-seq, reconstruction of 16 sections of Drosophila embryo individual measured by Stereo-seq, and reconstruction of a complete Mouse brain measured by BARseq, including 40 sections with sectioning interval as far as 200 μm. Among the methods, registered result by ST-GEARS exhibited the highest intra-structural consistency measured by MSSIM for two hippocampus sections separated by a single layer of neurons. On 16 sections of a Drosophila embryo individual, our method’s outstanding accuracy is indicated by both MSSIM and smoothness of tissue area changes. Importantly, ST-GEARS provides more reliable embryo morphology, precise tissue regionalization, and accurate marker gene distribution under hybridization evidence compared to existing approaches. This suggests that ST-GEARS provides higher quality tissues, cells, and genes information. On mouse brain sections with large intervals of 200 μm, ST-GEARS avoided positional and angular misalignments that occur in result of PASTE and PASTE2. The improvement was quantified by a higher MSSIM. Both hemisphere morphology and cortex layer regionalization were reflected in the result of 3D reconstruction by ST-GEARS. The successful representation of important structural and functional features in the aforementioned studies collectively underscores ST-GEARS’ reliability and capability for advancing 3D downstream research, enabling more comprehensive and insightful analysis of complex biological systems.

To further enhance and extend our method, opportunities in various aspects are anticipated to be explored. Firstly, tasks aimed at improving data preprocessing, including but not limited to batch effect removal and diffusion correction, are expected to be integrated into our method, considering their coupling property with registration task itself: inaccuracies in input data introduce perturbations to anchors optimization, while recovered spatial information of our method may assist data quality enhancement by providing registered sections. Secondly, the ST-GEARS’ Distributive Constraint takes rough grouping information as its input, which may potentially introduce computational burden during the reconstruction process. To address this, an automatic step is expected to be developed to reliably cluster spots while maintaining computational efficiency of the overall process. This step can be integrated into our method either as preprocessing, or as a coupling task, similarly to our expectation of data quality enhancement. Finally, we envision incorporating a wider scope of anchors applications into our existing framework. such as information integration of sections across time, across modalities and even across species. With interpretability, robustness and accuracy provided by ST-GEARS, we anticipate its applications and extension in various areas of biological and medical research. We believe that our method can help address a multitude of questions regarding growth and development, disease mechanisms, and evolutionary processes.

## Methods

### FGW OT Description

We introduce 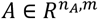 to describe normalized count of unique molecular identifiers (UMIs) of expressed genes on section A, where *n*_*A*_ denotes number of spots in slice A, and *m* denotes number of genes that are captured in both sections. Similarly, we describe gene expression on section B as 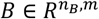, with genes arranged in the same order as in *A*. We introduce 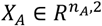 to describe locations of spots in section A, with spots arranged in the same order as in rows of matrix *A*; and similarly 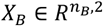, including spots locations in section B, arranged in the same order as in matrix *B*.

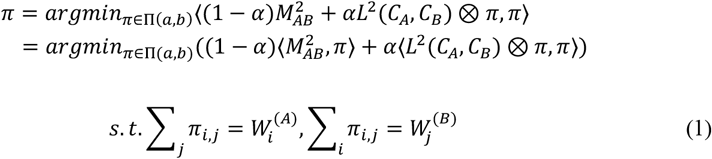

where 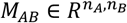 describes the similarity of each pair of spots respectively located on section A and B, formulated as 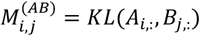, with *KL* denoting Kullback-Leibler (KL) divergence[46]. KL divergence measures gene expression difference between section A and B, taking the expression level of different genes as a probability distribution. 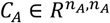 describes spatial structure of section A using intra spot pairwise Euclidean distances, with 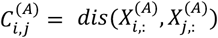, and *dis* denotes Euclidean distance measure of spots location. 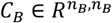 describes spatial structure of section B, with 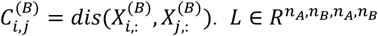 defines the similarity between all pairwise distances within slice A and slice B, with *l* 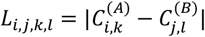. ⊗ denotes Kronecker product of two matrices; ⟨, ⟩ denotes matrix multiplication. 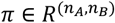 stores weighted anchors between spots from the two sections, with row and column index specifying the identity of spots, and element value representing probability which equals to weight of the anchor joining them, with 0 meaning non-existence of anchors. With 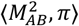, the similarity of joined spots are measured. With ⟨*L*^2^(*C*_*A*_, *C*_*B*_) ⊗ *π, π*⟩, similarity between all spots pair distances in section A, and pair distance of their anchoring spots in section B, is measured. ⟨*L*^2^(*C*_*A*_, *C*_*B*_) ⊗ *π, π*⟩ describes similarity between spatial structures under the anchors’ connection. *α* ∈ [0,1] denotes regularization factor, which specifies the relative importance of structure similarity compared to expression similarity, in the optimization process.

### Distributive Constraints

We introduce 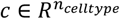, with each element specifying a cell type that appeared in both section A and B, and *n*_*celltype*_ denoting number of cell types. We use 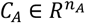 and 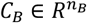 to describe the annotated cell type of each spot in section A and B, respectively.

First, for each cell type *c*_*i*_, find out its sum of expression matrix averaged by spots:

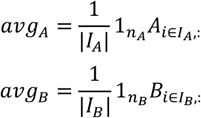

where both 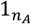 and 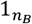 are defined as row vectors of ones:

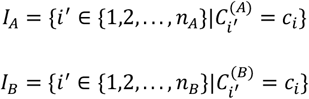

Next, for each cell type *c*_*i*_, measure the distance, described by KL divergence, of average expression matrix across sections, then map the distance by logistic kernel, to emphasize differences between highly consistent cell types.

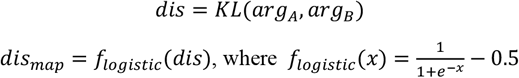

Finally, transform distance measure of all cell types to similarity measure, map them back to each spot on both sections and apply normalization on the result. ( 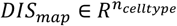 describes mapped distance of all cell types in *c*.)

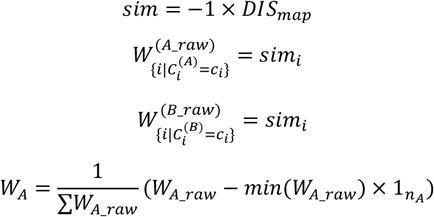

where 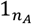 is a row vector of ones.

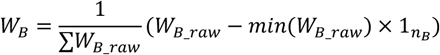

where 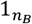 is a row vector of ones.

### Self-adaptive Regularization

In FGW OT formulation, a regularization factor was included to specify the relative importance of structural similarity compared to expression similarity during optimization. Our method includes a Self-adaptive Regularization method that determines the factor value, that induces highest overall accuracy of anchors despite of varying situations including but not limited to section distances, spot sizes, extent of distortions, and data quality such as level of diffusion. Our method respectively adopts factors on multiple scales including 0.8, 0.4, 0.2, 0.1, 0.05, 0.025, 0.013, and 0.006. The candidate values vary exponentially, for ST-GEARS to find the optimal term regardless of scale differences between expression and structural term in (1). The accuracy of each set of optimized anchors by every regularization factor was evaluated, by measuring weighted percentage 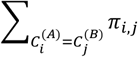 of anchors that join spots with same cell types. The regularization factor value that achieves highest accuracy is then adopted by our method.

### Elastic Field Inference

#### Finding spots with highest probability

With sections sequenced in cross-sectioning order, our method determines the most adjacent section(s) for each section, on both of its anterior and posterior sides. Exceptionally, if a section is on anterior or posterior end of a tissue, only one neighboring section is to be determined. For each section with *N* spots, we calculate 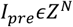, which stores the indices of mapped spots with highest probability, on its section on anterior side; as well as 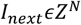, which stores the indices of mapped spots with highest probability, on its neighboring section on posterior side. Notably, not every spot in a selected section has its own anchored spot, due to multiple strategies including distributive constraint and anchors filtration, hence their corresponding element in *I*_*pre*_ and *I*_*next*_ are null.

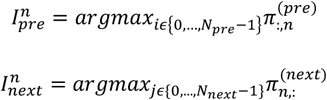

Thereinto, *π*_*pre*_ denotes the weighted anchors between the section and its anterior section, while *π*_*next*_ denotes the weighted anchors between the section and its posterior section.

Next, our method calculates location differences of spots on current section, and their mapped spots on anterior and posterior sections:

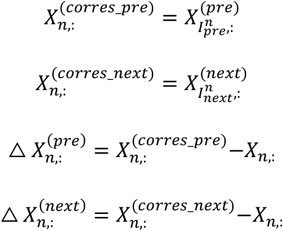

where *X* ∈ *R*^*N*,2^ denotes spots location of current section after rigid registration, and *X*_*pre*_ ∈ *R*^*N*,2^ and *X*_*next*_ ∈ *R*^*N*,2^ denote spots location of anterior and posterior section after rigid alignment, respectively.

#### Elastic Field establishment

Our method transforms the location differences from the form of individual values to displacement fields, with grid location indicating spot location, and grid value representing location differences between current spot, and its mapped spots on adjacent sections.

The displacement field is a matrix with height of *H* and width of *W*, where

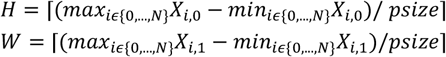

with *psize* representing distance between adjacent spot centers.

Spot location after rigid registration is pixelized, to correspond to its location in the field:

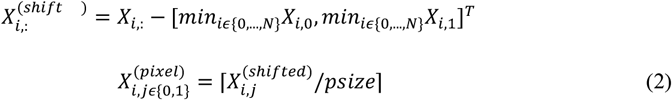

By the end of Eq (2), mapping between pixel locations, and their displacement values are established. To fill in every grid of the displacement field, 2d nearest interpolation method[47] was adopted, which fills in every grid which does not have corresponding spot or connection, with the displacement of its neighboring spot. Eventually, four fields without empty values are generated for each section, including *F*^(x_pre)^ and *F*^(y_pre)^ that denotes Elastic Field between anterior section and current section, respectively on horizontal and vertical direction; and *F*^(x_next)^ and *F*^(y_next)^ denoting Elastic Field between posterior section and current section on the same two directions:

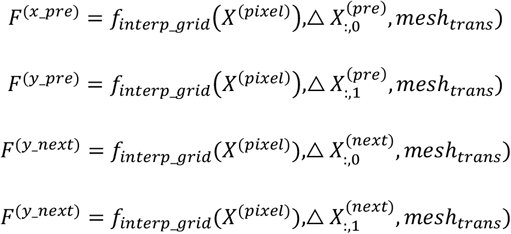

thereinto 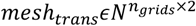 denotes grid coordinates of the designed field, with *n*_*grids*_ = *H* × *W*. And *f*_*interp_grid*_ denotes the nearest interpolation method.

#### 2D Gaussian Denoising

The computed field describes displacement distribution of each current section’s adjacent section(s) compared to its own. As the exerted force which causes the displacement passes through the whole section of tissue, the field value is expected to change smoothly across different locations[48][49][50]. Our method makes use of this property, to reduce errors in the field induced by both raw data defection and inaccuracies caused by upper stream algorithms. Gaussian filtering[51][52] is adopted to implement this strategy similarly to image denoising processes[53][54], which filters out dispersive outliers by replacing each grid value with the weighted average of its neighboring area:

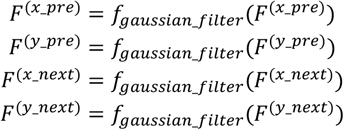

where *f*_*gaussian_filter*_ denotes the method of Gaussian filtering.

### Bi-sectional Fields Application

#### Bi-sectional Fields Application Plan

Our method applies the displacement field to each rigidly aligned sections to achieve elastic registration. To attain a distortion correction plan for each section, the displacement of both anterior and posterior sections compared to its own is computed, by retrieving offsets value from each displacement field:

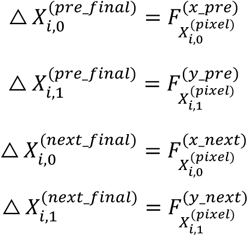

Notice that the first section in sequenced slices is only equipped with △ *X*_*next_final*_, while the last section is only equipped with △ *X*_*pre_final*_.

To correct the distortion, a correction plan by averaging displacement field from both anterior and posterior sections, is applied onto the coordinates of spots after rigid alignment. We introduce *X*_*final*_ ∈ *R*^*N*,2^ to represent recovered spatial information through elastic registration, and

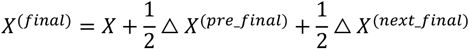

In the first section in sequenced slices,

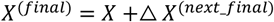

In the last section in sequenced slices,

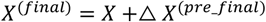

The validity of this plan is proved as following.

#### Proof of validity of Bi-sectional Fields Application

Take section A, B, and C as an example of a sequence of sections, with *X*_*A*_, *X*_*B*_ and *X*_*C*_ denoting their spots’ spatial information after rigid alignment, with *X*_*A_insitu*_, *X*_*B_insitu*_ and *X*_*C_insitu*_ denoting their *in situ* spatial information. We model the distortion occurred to the slices as changes on their spatial information, written as *X*_*A_dis*_, *X*_*B_dis*_ and *X*_*C_dis*_.

The corrected spatial information,

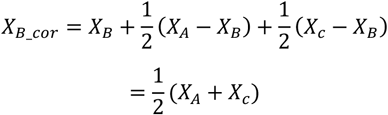

Thereinto,

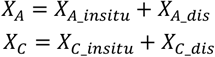

Hence,

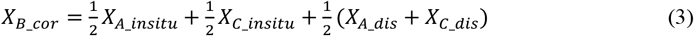

Based on the *in situ* morphological consistency across sections, spatial information of section B can be approximated by an average of information of A and C, written as

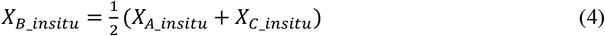

Given that *X*_*A_dis*_ and *X*_*C_dis*_ can be seen as independent and identically distributed sets of variables,

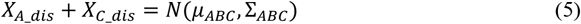

where *µ*_*ABC*_ is the universal mean, and Σ_*ABC*_ is the variance of the 2d displacement information.

Inserting the terms (4) and (5) back to Eq (3) gives

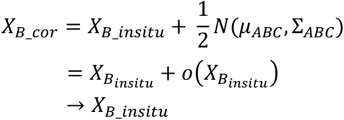

indicating the proximity of corrected spatial information to *in situ* spatial information.

#### Evaluation Metrix

We evaluated the accuracy of anchors by index of Mapping Accuracy, and measured the reconstruction effect by MSSIM and SI-STD-DI, in both elastic effect study and overall methodology comparison.

#### Mapping Accuracy

Designed and adopted by PASTE[27], Mapping Accuracy calculates the weighted percentage of anchors joining spots with same annotation.

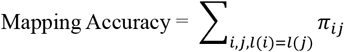

#### MSSIM index

MSSIM measures the accuracy of registration, based on the assumption that in some sectioning positions, tissue morphology remains almost consistent across slices. The method quantifies the accuracy, by measuring the similarity of cell type distribution of such section pairs.

To implement the quantification, first, structurally consistent section pairs are selected among all sections arranged in sequence.

Next, on each section from the pair, transformation from individual spots to a complete image is implemented, by gridding the rectangular area that surrounds the tissue, and assigning each grid of a value that represents the cell type which occurs most frequently in the grid. The resulted image describes the cell type distribution of the section.

Finally, similarity between each pair of images is measured, by index of MSSIM[55]. The method generates a window with fixed size, slides the window simultaneously on both images, and compares the two framed parts by windows on their intensity, contrast, and structures. Among those, the intensity difference is measured by difference of average pixel values, the contrast difference is measured by comparing variance of the two sets of framed pixel values, and the structure difference is measured by comparing their covariances. A Structural Similarity of Images (SSIM) index is calculated for each position of the window using 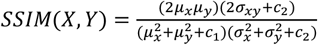,where *µ*_*x*_ and *µ*_*y*_ denote average pixel values of the frames, *σ*_*x*_ and *σ*_*y*_ denote variances of the frames, and *σ*_*xy*_ denotes covariances of the two frames. *c*_1_ and *c*_2_ are constants to avoid 0 value of the divisor. Averaging the SSIM value across all windows gives the final MSSIM result of the two sections.

#### SI-STD-DI

SI-STD-DI measures smoothness of area changing across sections along a fixed axis, by calculating the standard deviation of area changes on each pair of adjacent sections and scale the result by dividing it by average area.

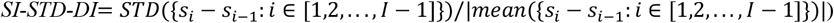

## Dataset availability

All data used in this research were collected from published sources. DLPFC data was obtained from the research: Transcriptome-scale Spatial Gene Expression in the Human Dorsolateral Prefrontal Cortex, with data downloading link of http://research.libd.org/spatialLIBD/index.html; Drosophila embryo and Drosophila larva data were collected from High-resolution 3d Spatiotemporal Transcriptomic Maps of Developing Drosophila Embryos and Larvae, with the dataset link of https://db.cngb.org/stomics/datasets/STDS0000060. Mouse brain data was collected from research: Modular cell type organization of cortical areas revealed by in situ sequencing. The download link is: https://data.mendeley.com/datasets/8bhhk7c5n9/draft?a=f37296fe-dc00-46a6-49e69-46f4c3d0deec. All datasets were generated on Spatial Transcriptomics platform, with DLPFC data generated by Visium technology of 10x Genomics, Mouse brain data generated by BARseq of Cold Spring Harbor Laboratory, while Drosophila embryo and larva generated by Stereo-seq technology of BGI.

## Code Availability

The methods of ST-GEARS is packaged, and distributed as an open-source, publicly available repository: https://github.com/STOmics/ST-GEARS.

## Acknowledgements

This work is part of the “SpatioTemporal Omics Consortium” (STOC) paper package. A list of STOC members is available at: http://sto-consortium.org. We acknowledge the Stomics Cloud platform (https://cloud.stomics.tech/) for providing convenient ways for analyzing spatial omics datasets. We thank Weizhen Xue for the inspirational discussion towards design of Distributive Constraints. We thank Yating Ren for her advice towards a more efficient code implementation. We thank Dr. Xiaojie Qiu and Dr. Yinqi Bai for the discussion on the registration topic and their advice on our work. We acknowledge the CNGB Nucleotide Sequence Archive (CNSA) of China National GeneBank DataBase (CNGBdb) for maintaining the Drosophila database.

## Author contributions

Tianyi Xia was responsible of method design, analysis design and implementation, as well as drafting of this manuscript. Dr. Luni Hu participated in structure design of the applications. Lulu Zuo was in part of 3D visualizations design, and she helps maintain our online repository. Dr. Yunjia Zhang provided insights in anchors results interpretation of DLPFC dataset, and in accuracy analysis of mouse brain dataset. Dr. Mengyang Xu revised this article. Lei Zhang and Bowen Ma offered numerous suggestions to enhance computational efficiency, in both memory and time. Taotao Pan and Chuan Chen provided suggestions in data preprocessing. Qin Lu, Lei Cao, Bohan Zhang, Junfu Guo, Chang Shi and Mei Li provided suggestions for this study. Dr. Shuangsang Fang supervised this study in structure and analysis design, and she revised this article. Chao Liu, Yuxiang Li and Yong Zhang supervised this study.

## References

[1]. Marx, V.: Method of the year: spatially resolved transcriptomics. Nature methods 18(1), 9–14 (2021).

[2]. Yue, L., Liu, F., Hu, J., Yang, P., Wang, Y., Dong, J., Shu, W., Huang, X., Wang, S.: A guidebook of spatial transcriptomic technologies, data resources and analysis approaches. Computational and Structural Biotechnology Journal 21 (2023)

[3]. Park, H.-E., Jo, S.H., Lee, R.H., Macks, C.P., Ku, T., Park, J., Lee, C.W., Hur, J.K., Sohn, C.H.: Spatial transcriptomics: Technical aspects of recent developments and their applications in neuroscience and cancer research. Advanced Science 10(16), 2206939 (2023)

[4]. Gyllborg, D., Langseth, C.M., Qian, X., Choi, E., Salas, S.M., Hilscher, M.M., Lein, E.S., Nilsson, M.: Hybridization-based in situ sequencing (hybiss) for spatially resolved transcriptomics in human and mouse brain tissue. Nucleic acids research 48(19), 112–112 (2020).

[5]. Chen, X., Sun, Y.-C., Zhan, H., Kebschull, J.M., Fischer, S., Matho, K., Huang, Z.J., Gillis, J., Zador, A.M.: High-throughput mapping of long-range neuronal projection using in situ sequencing. Cell 179(3), 772–786 (2019).

[6]. Wang, X., Allen, W.E., Wright, M.A., Sylwestrak, E.L., Samusik, N., Vesuna, S., Evans, K., Liu, C., Ramakrishnan, C., Liu, J., et al.: Three-dimensional intact-tissue sequencing of single-cell transcriptional states. Science 361(6400), 5691(2018).

[7]. Qin, D.: Next-generation sequencing and its clinical application. Cancer biology & medicine 16(1), 4 (2019).

[8]. Chen, A., Liao, S., Cheng, M., Ma, K., Wu, L., Lai, Y., Yang, J., Li, W., Xu, J., Hao, S., et al.: Large field of view-spatially resolved transcriptomics at nanoscale resolution. BioRxiv 2021 (2021).

[9]. Stickels, R.R., Murray, E., Kumar, P., Li, J., Marshall, J.L., Di Bella, D.J., Arlotta, P., Macosko, E.Z., Chen, F.: Highly sensitive spatial transcriptomics at near-cellular resolution with slide-seqv2. Nature biotechnology 39(3), 313–319 (2021).

[10]. Moses, L., Pachter, L.: Museum of spatial transcriptomics. Nature Methods 19(5), 534–546 (2022)

[11]. Moor, A.E., Itzkovitz, S.: Spatial transcriptomics: paving the way for tissue-level systems biology. Current opinion in biotechnology 46, 126–133 (2017).

[12]. Zhou, R., Yang, G., Zhang, Y., Wang, Y.: Spatial transcriptomics in development and disease. Molecular Biomedicine 4(1), 32 (2023).

[13]. Li, Z., Peng, G.: Spatial transcriptomics: new dimension of understanding biological complexity. Biophysics Reports 8(3), 119 (2022).

[14]. Williams, C.G., Lee, H.J., Asatsuma, T., Vento-Tormo, R., Haque, A.: An introduction to spatial transcriptomics for biomedical research. Genome Medicine 14(1), 1–18 (2022).

[15]. Walker, B.L., Cang, Z., Ren, H., Bourgain-Chang, E., Nie, Q.: Deciphering tissue structure and function using spatial transcriptomics. Communications biology 5(1), 220 (2022).

[16]. Atta, L., Fan, J.: Computational challenges and opportunities in spatially resolved transcriptomic data analysis. Nature Communications 12(1), 5283 (2021).

[17]. Velten, B., Braunger, J.M., Argelaguet, R., Arnol, D., Wirbel, J., Bredikhin, D., Zeller, G., Stegle, O.: Identifying temporal and spatial patterns of variation from multimodal data usingmefisto. Nature methods 19(2), 179–186 (2022).

[18]. Townes, F.W., Engelhardt, B.E.: Nonnegative spatial factorization applied to spatial genomics. Nature Methods 20(2), 229–238 (2023).

[19]. Verma, A., Engelhardt, B.: A bayesian nonparametric semi-supervised model for integration of multiple single-cell experiments. bioRxiv, 2020–01 (2020).

[20]. Svensson, V., Teichmann, S.A., Stegle, O.: Spatialde: identification of spatially variable genes. Nature methods 15(5), 343–346 (2018).

[21]. Dries, R., Zhu, Q., Dong, R., Eng, C.-H.L., Li, H., Liu, K., Fu, Y., Zhao, T., Sarkar, A., Bao, F., et al.: Giotto: a toolbox for integrative analysis and visualization of spatial expression data. Genome biology 22, 1–31 (2021).

[22]. Wang, M., Hu, Q., Lv, T., Wang, Y., Lan, Q., Xiang, R., Tu, Z., Wei, Y., Han, K., Shi, C., et al.: High-resolution 3d spatiotemporal transcriptomic maps of developing drosophila embryos and larvae. Developmental Cell 57(10), 1271–1283(2022).

[23]. Mohenska, M., Tan, N.M., Tokolyi, A., Furtado, M.B., Costa, M.W., Perry, A.J., Hatwell-Humble, J., Duijvenboden, K., Nim, H.T., Ji, Y.M., et al.: 3d-cardiomics: a spatial transcriptional atlas of the mammalian heart. Journal of molecular and cellular cardiology 163, 20–32 (2022)

[24]. Vickovic, S., Schapiro, D., Carlberg, K., Lötstedt, B., Larsson, L., Hildebrandt, F., Korotkova, M., Hensvold, A.H., Catrina, A.I., Sorger, P.K., et al.: Three-dimensional spatial transcriptomics uncovers cell type localizations in the human rheumatoid arthritis synovium. Communications Biology 5(1), 129 (2022)

[25]. Rao, A., Barkley, D., Fran ça, G.S., Yanai, I.: Exploring tissue architecture using spatial transcriptomics. Nature 596(7871), 211–220 (2021).

[26]. Bergenstråhle, J., Larsson, L., Lundeberg, J.: Seamless integration of image and molecular analysis for spatial transcriptomics workflows. BMC genomics 21(1), 1–7 (2020).

[27]. Zeira, R., Land, M., Strzalkowski, A., Raphael, B.J.: Alignment and integration of spatial transcriptomics data. Nature Methods 19(5), 567–575 (2022).

[28]. Liu, X., Zeira, R., Raphael, B.: Paste2: Partial alignment of multi-slice spatially resolved transcriptomics data. In: Research in Computational Molecular Biology: 27th Annual International Conference, RECOMB 2023, Istanbul, Turkey, April 16–19, 2023, Proceedings, vol. 13976, p. 210 (2023). Springer Nature

[29]. Jones, A., Townes, F.W., Li, D., Engelhardt, B.E.: Alignment of spatial genomics data using deep gaussian processes. Nature Methods, 20(9), 1379–1387, (2023).

[30]. Xia, C.-R., Cao, Z.-J., Tu, X.-M., Gao, G.: Spatial-linked alignment tool (slat) for aligning heterogenous slices properly. bioRxiv, 2023–04 (2023).

[31]. Qiu, X., Zhu, D.Y., Yao, J., Jing, Z., Zuo, L., Wang, M., Min, K.H., Pan, H., Wang, S., Liao, S., et al.: Spateo: multidimensional spatiotemporal modeling of single-cell spatial transcriptomics. BioRxiv, 2022–12 (2022).

[32]. Guo, L., Li, Y., Qi, Y., Huang, Z., Han, K., Liu, X., Liu, X., Xu, M., Fan, G.: Vt3d: a visualization toolbox for 3d transcriptomic data. Journal of Genetics and Genomics (2023).

[33]. Fang, S., Xu, M., Cao, L., Liu, X., Bezulj, M., Tan, L., Yuan, Z., Li, Y., Xia, T., Guo, L., et al.: Stereopy: modeling comparative and spatiotemporal cellular heterogeneity via multi-sample spatial transcriptomics. bioRxiv, 2023–12 (2023).

[34]. Titouan, V., Courty, N., Tavenard, R., Flamary, R.: Optimal transport for structured data with application on graphs. In: International Conference on Machine Learning, pp. 6275–6284 (2019). PMLR.

[35]. Maynard, K.R., Collado-Torres, L., Weber, L.M., Uytingco, C., Barry, B.K., Williams, S.R., Catallini, J.L., Tran, M.N., Besich, Z., Tippani, M., et al.: Transcriptome-scale spatial gene expression in the human dorsolateral prefrontal cortex. Nature neuroscience 24(3), 425–436 (2021).

[36]. Rodriques, S.G., Stickels, R.R., Goeva, A., Martin, C.A., Murray, E., Vanderburg, C.R., Welch, J., Chen, L.M., Chen, F., Macosko, E.Z.: Slide-seq: A scalable technology for measuring genome-wide expression at high spatial resolution. Science 363(6434), 1463–1467 (2019).

[37]. Chen, X., Fischer, S., Zhang, A., Gillis, J., Zador, A.: Modular cell type organization of cortical areas revealed by in situ sequencing. BioRxiv, 2022–11(2022).

[38]. Abdolhosseini, F., Azarkhalili, B., Maazallahi, A., Kamal, A., Motahari, S.A., Sharifi-Zarchi, A., Chitsaz, H.: Cell identity codes: understanding cell identity from gene expression profiles using deep neural networks. Scientific reports 9(1), 2342 (2019).

[39]. Efroni, I., Ip, P.-L., Nawy, T., Mello, A., Birnbaum, K.D.: Quantification of cell identity from single-cell gene expression profiles. Genome biology 16, 1–12 (2015).

[40]. Lacoste-Julien, S.: Convergence rate of frank-wolfe for non-convex objectives. arXiv preprint arXiv:1607.00345 (2016).

[41]. Wahba, G.: A least squares estimate of satellite attitude. SIAM review 7(3), 409–409 (1965).

[42]. Williams, C.G., Lee, H.J., Asatsuma, T., Vento-Tormo, R., Haque, A.: An introduction to spatial transcriptomics for biomedical research. Genome Medicine 14(1), 1–18 (2022).

[43]. Clifton, K., Anant, M., Aimiuwu, O.K., Kebschull, J.M., Miller, M.I., Tward, D., Fan, J.: Alignment of spatial transcriptomics data using diffeomorphic metric mapping. bioRxiv, 2023–04 (2023).

[44]. Lanjakornsiripan, D., Pior, B.-J., Kawaguchi, D., Furutachi, S., Tahara, T., Kat-suyama, Y., Suzuki, Y., Fukazawa, Y., Gotoh, Y.: Layer-specific morphological and molecular differences in neocortical astrocytes and their dependence on neuronal layers. Nature communications 9(1), 1623 (2018).

[45]. Benavides-Piccione, R., Regalado-Reyes, M., Fernaud-Espinosa, I., Kastanauskaite, A., Tapia-Gonz ález, S., Leéon-Espinosa, G., Rojo, C., Insausti, R., Segev, I., DeFelipe, J.: Differential structure of hippocampal ca1 pyramidal neurons in the human and mouse. Cerebral cortex 30(2), 730–752 (2020).

## References

[46]. Csiszár, I.: I-divergence geometry of probability distributions and minimization problems. The annals of probability, 146–158 (1975).

[47]. Schoenberg, I.J.: Contributions to the problem of approximation of equidistant data by analytic functions: Part a.—on the problem of smoothing or graduation. a first class of analytic approximation formulae. IJ Schoenberg Selected Papers, 3–57 (1988).

[48]. Zhou, H., Jayender, J.: Smooth deformation field-based mismatch removal in real-time. arXiv preprint arXiv:2007.08553 (2020).

[49]. Li, X., Hu, Z.: Rejecting mismatches by correspondence function. International Journal of Computer Vision 89, 1–17 (2010).

[50]. Li, X., Larson, M., Hanjalic, A.: Pairwise geometric matching for large-scale object retrieval. In: Proceedings of the IEEE Conference on Computer Vision and Pattern Recognition, pp. 5153–5161 (2015)

[51]. Bergholm, F.: Edge focusing. IEEE transactions on pattern analysis and machine intelligence (6), 726–741 (1987).

[52]. Marr, D., Hildreth, E.: Theory of edge detection. Proceedings of the Royal Society of London. Series B. Biological Sciences 207(1167), 187–217 (1980)

[53]. Mafi, M., Martin, H., Cabrerizo, M., Andrian, J., Barreto, A., Adjouadi, M.: A comprehensive survey on impulse and gaussian denoising filters for digital images. Signal Processing 157, 236–260 (2019).

[54]. Saxena, C., Kourav, D.: Noises and image denoising techniques: a brief survey. International journal of Emerging Technology and advanced Engineering 4(3), 14878–885 (2014).

[55]. Wang, Z., Bovik, A.C., Sheikh, H.R., Simoncelli, E.P.: Image quality assessment: from error visibility to structural similarity. IEEE transactions on image processing 13(4), 600–612 (2004).

