## Supplementary Materials for "ST-GEARS: Advancing 3D Downstream Research through Accurate Spatial Information Recovery"

### Supplementary Information

##### Data Preprocessing

Gene expression, cross sectional clustering or annotation, and spatial measuring of different spots across sections are required as input of ST-GEARS. Across different ST technologies, resolutions that could be as high as 500 nm can lead to over millions of input spots[8], causing computational burden in both time and memory. To deal with the situation, we include spots granularity adjusting is included as an optional preprocessing step, and we recommend users to turn on this option : when over 3000 spots are included in each section. In granularity adjusting, section area is gridded, with spots squared by each pixel summarized into one single spot. The strategy enables higher computational efficiency in both anchors computation and geospatial correction, without compromising accuracy of the reconstructed result. We applied this preprocessing onto Mouse brain dataset, summarizing each cell's MRNA expression into its belonging 200 μm wide pixels(Supplementary Fig. 19a).

For each summarized or raw spot, both gene expression and geospatial profile are utilized by ST-GEARS as its input. Unique molecular identifier (UMI) counts responsible for gene expression profile are linearly scaled to 0 to 10000, before log normalization.

**Reconstruct samples with different constraints settings**

We introduced Distributive Constraints in ST-GEARS to assign different levels of emphasis on different groups of spots, for an enhanced anchor and therefore reconstruction accuracy. We applied Distributive Constraints on registration and reconstruction of DLPFC, Drosophila embryo, Drosophila larva and Mouse hippocampus. However, on Mouse brain dataset, we did not adopt the setting, because a vast changing of cell types constitution was witnessed across sections with large distance of 200 μm. In such a circumstance, a distributive emphasis on spots according to cell types were not expected to function equally well with other applications, since it impedes anchors generation on spots with relative similar expressions but assigned to different cell types. In our code repository, we also remained Distributive Constraints as an optional parameter, which is recommended to not to apply in cases of vast grouping changes, cluster changes across sections, or during absence of reliable grouping information. Users are also encouraged to check out registration results of both settings to determine whether to adopt Distributive Constraints.

**Benchmarking**

To implement benchmarking analysis on PASTE, PASTE2 and GPSA, datasets were preprocessed as according to the requirements of the respective methods. Spots granularity adjusting was applied onto the Mouse brain dataset as input of the three methods, to maintain the same data size and granularity with ST-GEARS. Parameters were set to default during the computation of the methods.

### Supplementary Figures

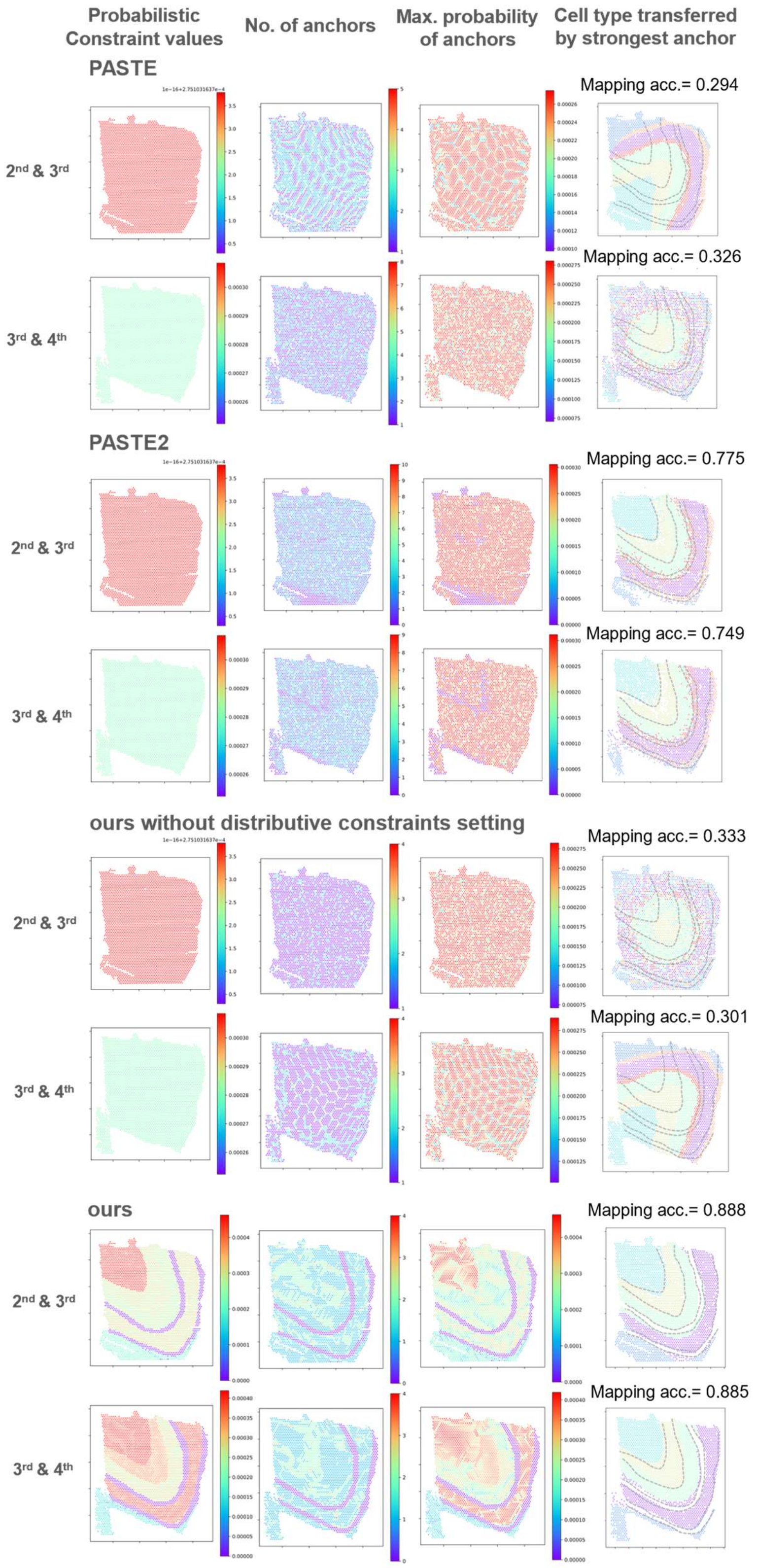


Supplementary Fig. 1: **Distributive emphasis of different cell types of the 2nd to 4th section of DLPFC causes the advanced anchor accuracy.** From the 1st to the 4th column are presented probabilistic constraints settings in problem formulating, no. of anchors computed on each spot, max. anchor probability value computed of each spot, and annotated cell types on the next sections mapped back to its previous sections through computed anchors, with mapping accuracy marked. The distinction of different cell types on the sections are marked by dotted lines. 1st and 2nd row show analysis results of PASTE, 3rd and 4th show results of PASTE2, 5th and 6th row show results of ST-GEARS without distributive constraints settings, and 7th and 8th rows show results of ST-GEARS with distributive constraints settings. In the results of each method, the upper row presents result of the 2nd and the 3rd sections, while the lower row presents results of the 3rd and 4th sections. As ST-GEARS adopts distributive constraints, it generates relatively more and higher probabilistic anchors on cell types with higher expressional consistency across sections, and hence it produces anchors with higher mapping accuracy.

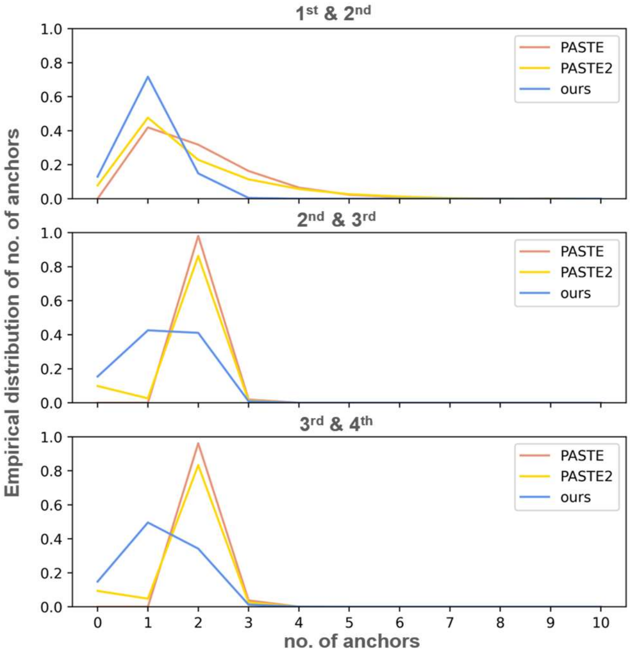


Supplementary Fig. 2: **Suppression of anchors generation on certain spots by ST-GEARS.** Shown here is the comparison of number of anchors distribution of DLPFC dataset computed by PASTE, PASTE2 and ST-GEARS. Each y value shows fraction of no. of anchors specified on x. Different from PASTE, certain percentage of spots have zero anchors generated by ST-GEARS.

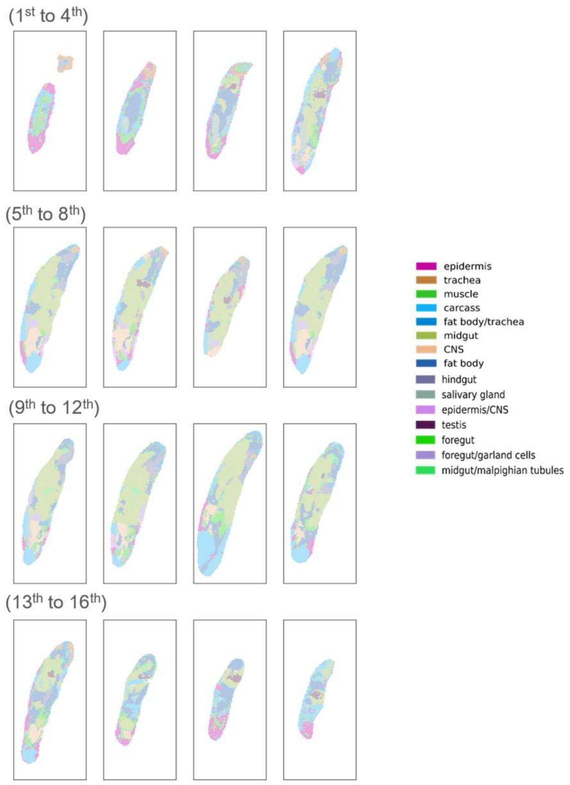


Supplementary Fig. 3: **Individual sections of Drosophila larva generated by rigid registration of ST-GEARS.**

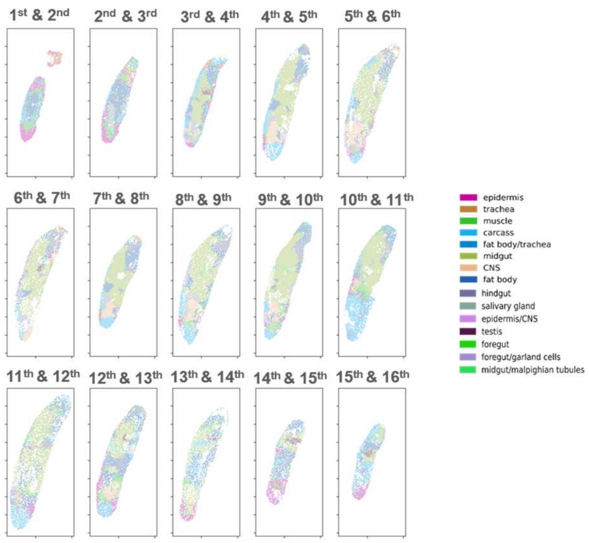


Supplementary Fig. 4: **Anchors correctly connect spots of Drosophila larva with same annotations.** The figure shows annotated cell types of next sections of Drosophila larva mapped back to their previous sections through computed anchors generated by ST-GEARS. The result corresponds well to original cell type distributions of the previous sections, indicating anchors connect spots with same annotations.

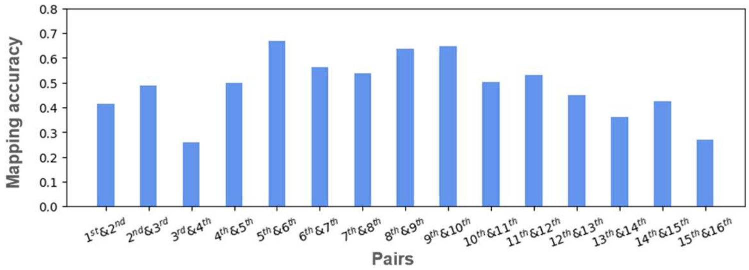


Supplementary Fig. 5: **Mapping accuracy of ST-GEARS on Drosophila larva.**

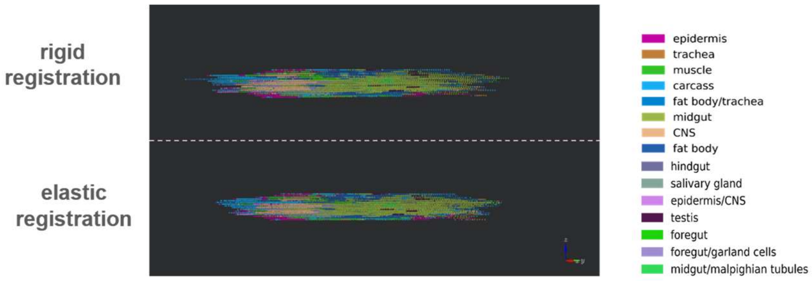


Supplementary Fig. 6: **Cell types of cross sections along Anterior-Posterior(A-P) of Drosophila larva.** Top row presents aligned results by rigid registration, and bottom row presents recovered coordinates by elastic registration, which is appended to rigid registration process.

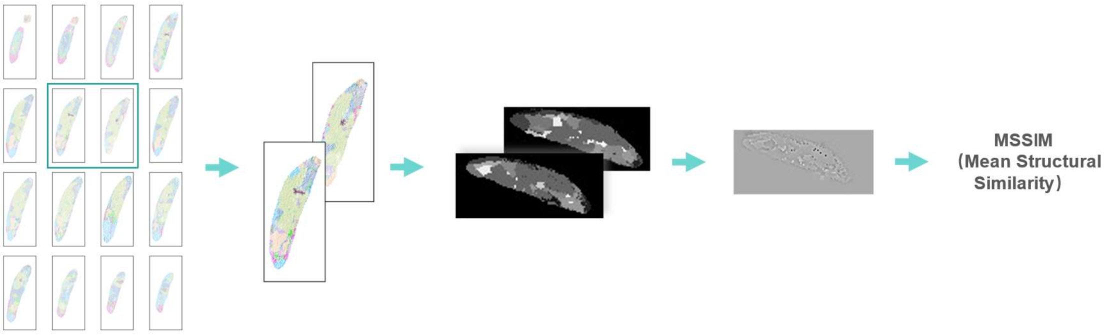


Supplementary Fig. 7: **Calculation of MSSIM index.** To score the registration result of multiple sections, structurally consistent pairs are picked up, and pixeled, with cell type that occur most times in each pixel representing the cell type of the pixel. Cell types are then transformed to grayscales, with different cell types taking different gray scales. Image MSSIM score of pairs is then calculated, representing MSSIM score of the registered pairs.

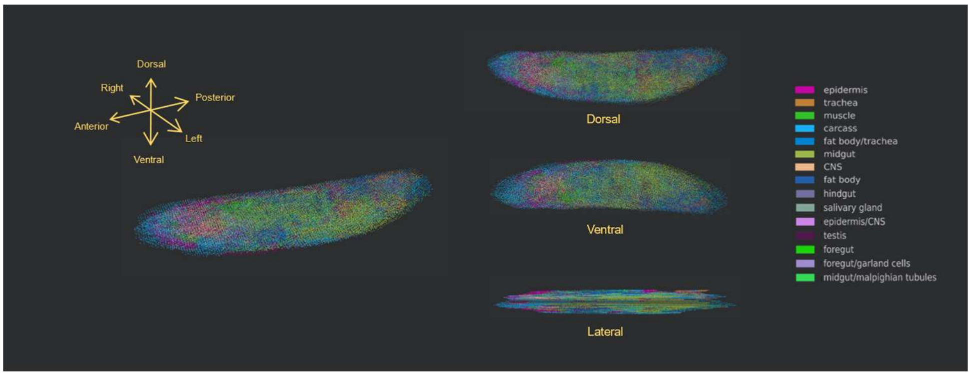


Supplementary Fig. 8: **Elastic process registers Drosophila sections by recovering its *in situ* geo-spatial profile.** Shown here are stacked sections of Drosophila larva generated by elastic registration appended to rigid registration of ST-GEARS, in perspective, dorsal, ventral, and lateral views.

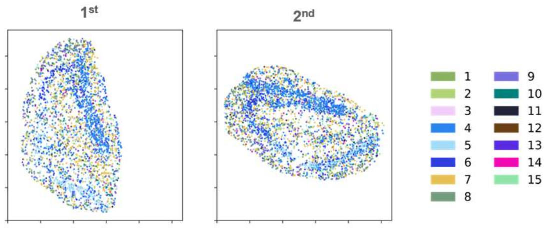


Supplementary Fig. 9: **Sagittal Mouse hippocampus sections before registration.**

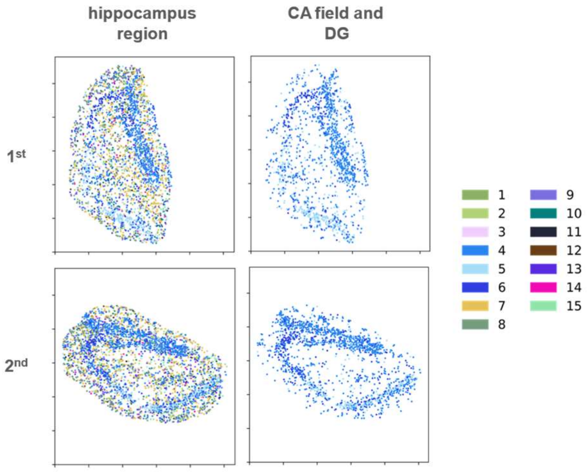


Supplementary Fig. 10: **Cornu Ammonis (CA) fields and Dentate Gyrus (DG) extraction of sagittal Mouse hippocampus sections.** Top row presents the 1st section of Mouse hippocampus, and bottom row represents the 2nd section. Left column presents all spots on the sections, while right column presents filtered spots that belong to CA fields and DG.

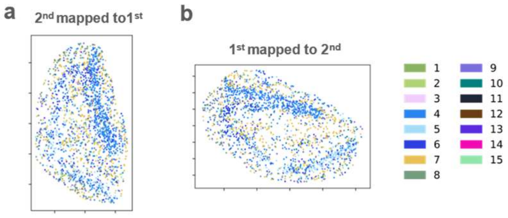


Supplementary Fig. 11: **Anchors correctly connect spots of Mouse hippocampus with same annotations. (a)** Annotated cell types of the 2nd section mapped back the 1st section through computed anchors generated by ST-GEARS. (b) Similar to (a), but cell types of the 1st section mapped to the 2nd. The result corresponds well to original cell type distributions of the sections, which means anchors connect spots with same annotations.

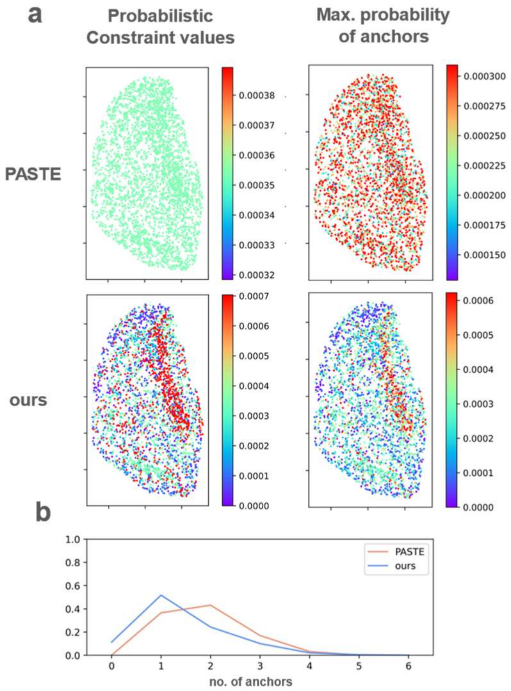


Supplementary Fig. 12: **Distributive emphasis of different cell types of Mouse hippocampus induces distributive anchors by ST-GEARS.** (a) In contrast to PASTE, ST-GEARS assigns distributive probabilistic constraints on different cell types of Mouse hippocampus as shown on the 1st row. By result, its generated maximum probabilities of spots' anchors are different across cell types, compared to PASTE, as shown on the 2nd row. **(b)** Different from PASTE, certain percentage of spots have zero anchors generated by ST-GEARS.

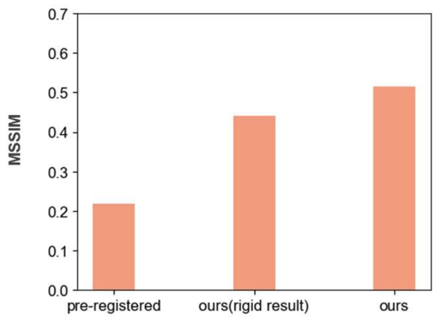


Supplementary Fig. 13: **Elastic registration further improves registration result of Mouse hippocampus based on rigid registration.** Shown here is the MSSIM index of pre-registered, rigid registered and elastic registered section pair of Mouse hippocampus.

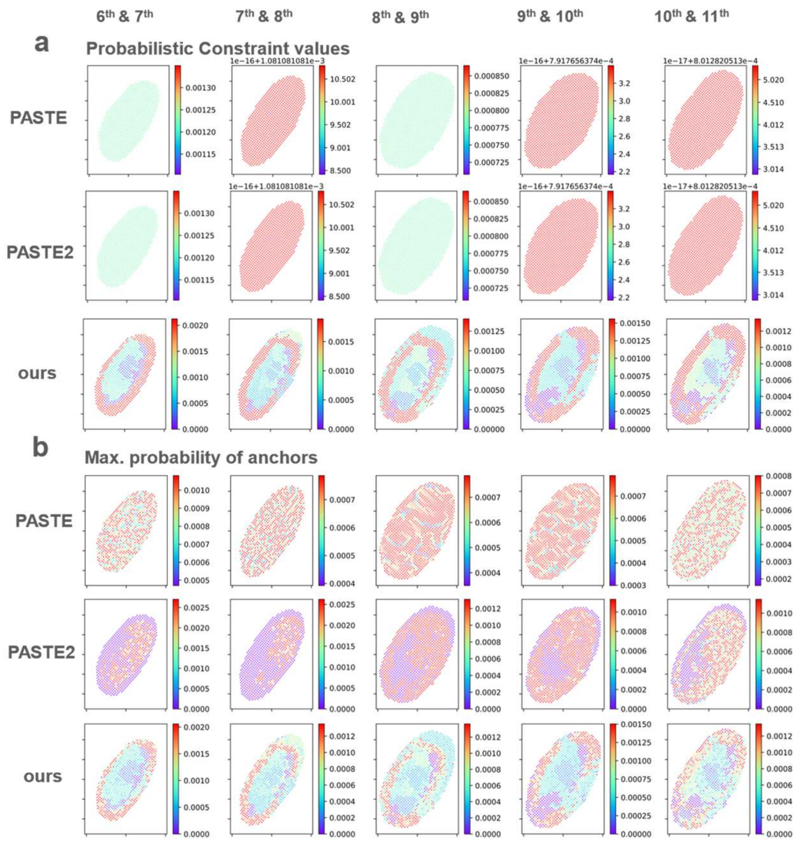


Supplementary Fig. 14: **Distributive emphasis of different cell types of Drosophila embryo induces distributive anchors by ST-GEARS.** (a) In contrast to PASTE and PASTE2, ST-GEARS assigns distributive probabilistic constraints on different cell types of Drosophila embryo as shown on the 1st , 2nd and 3rd row (b) By result, its generated maximum probabilities of spots' anchors are different across cell types, compared to PASTE and PASTE2, as shown on the 4th, 5th and 6th row.

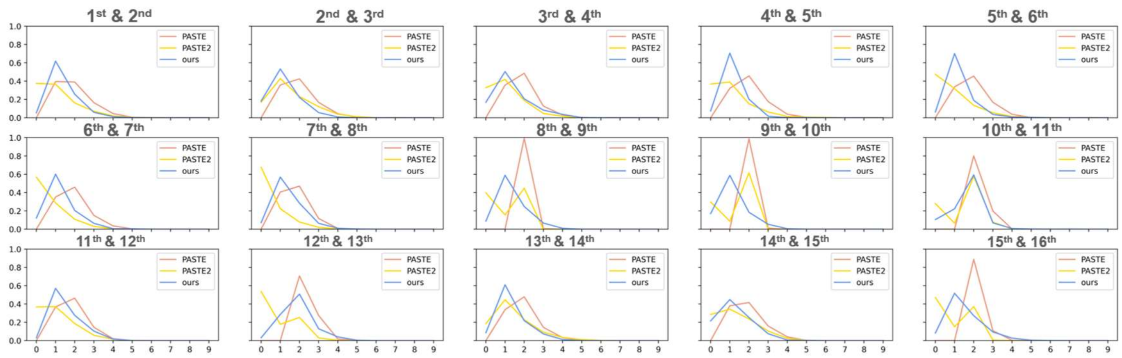

Supplementary Fig. 15: **Suppression of anchors generation on certain spots of Drosophila embryo by ST-GEARS.** Shown here is the comparison of number of anchors distribution of Drosophila embryo dataset computed by PASTE, PASTE2 and ST-GEARS. Each y value shows fraction of no. of anchors specified on x. Different from PASTE, certain percentage of spots have zero anchors generated by ST-GEARS.

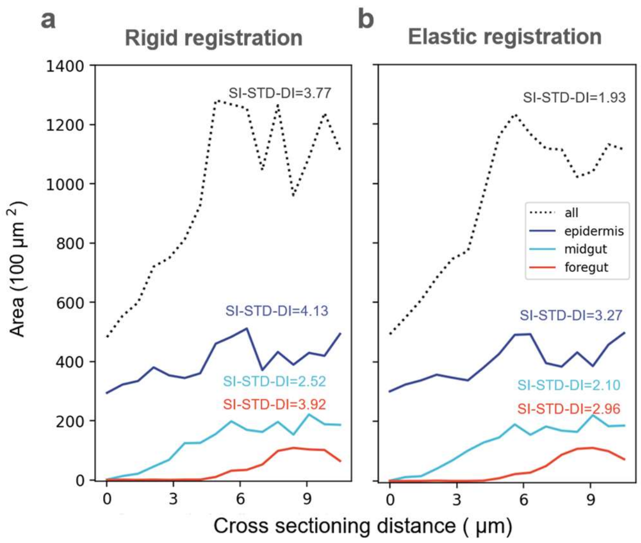


Supplementary Fig. 16: **Elastic registration in ST-GEARS smooths tissue shapes of Drosophila embryo.** (a) Shown here is area changes of epidermis, foregut, and midgut and overall body, along sectioning direction after rigid registration of ST-GEARS. Each area data is calculated based on pixelating the region of tissues, on each registered section, and summing up area of respective pixels. Scale-independent Standard Deviation of Differences (SSI_STD_DI) of each curve is calculated and marked as smoothness index of the curve. (b) Shown here is area changes of the 3 same tissues as (a) and overall body, along sectioning direction after elastic registration of ST-GEARS. Area changes are visually and quantitatively smoother after elastic registration than after rigid registration only.

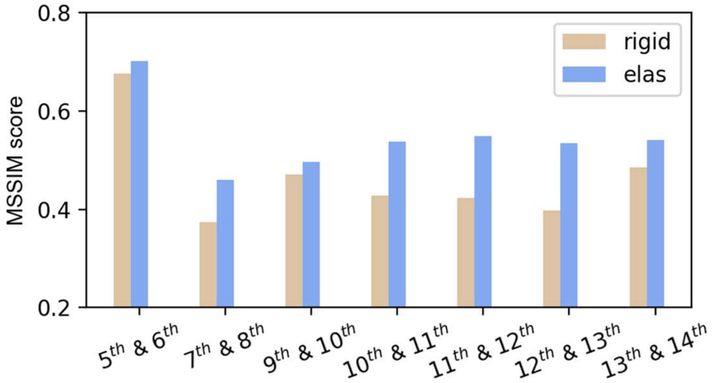


Supplementary Fig. 17: **Elastic registration in ST-GEARS enhances cross-sectional consistency in structurally consistent positions of Drosophila embryo.** Shown here **is** the comparison of Mean Structural Similarity (MSSIM) index of rigid and elastic registration result, of structurally consistent section pairs of Drosophila embryo. The similarity index is higher on elastic than on rigid result.

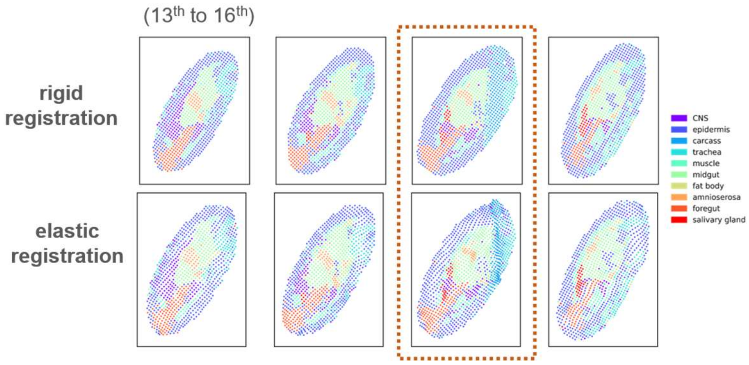


Supplementary Fig. 18: **Elastic registration fixes flaw of Drosophila embryo data.** Top row shows 13$^{th}$ to 16$^{th}$ sections after rigid registration, and bottom row shows corresponding sections after elastic registration. On the 15$^{th}$ section, the flaw region between 2 parallel lines caused during experimental phase was fixed by elastic registration.

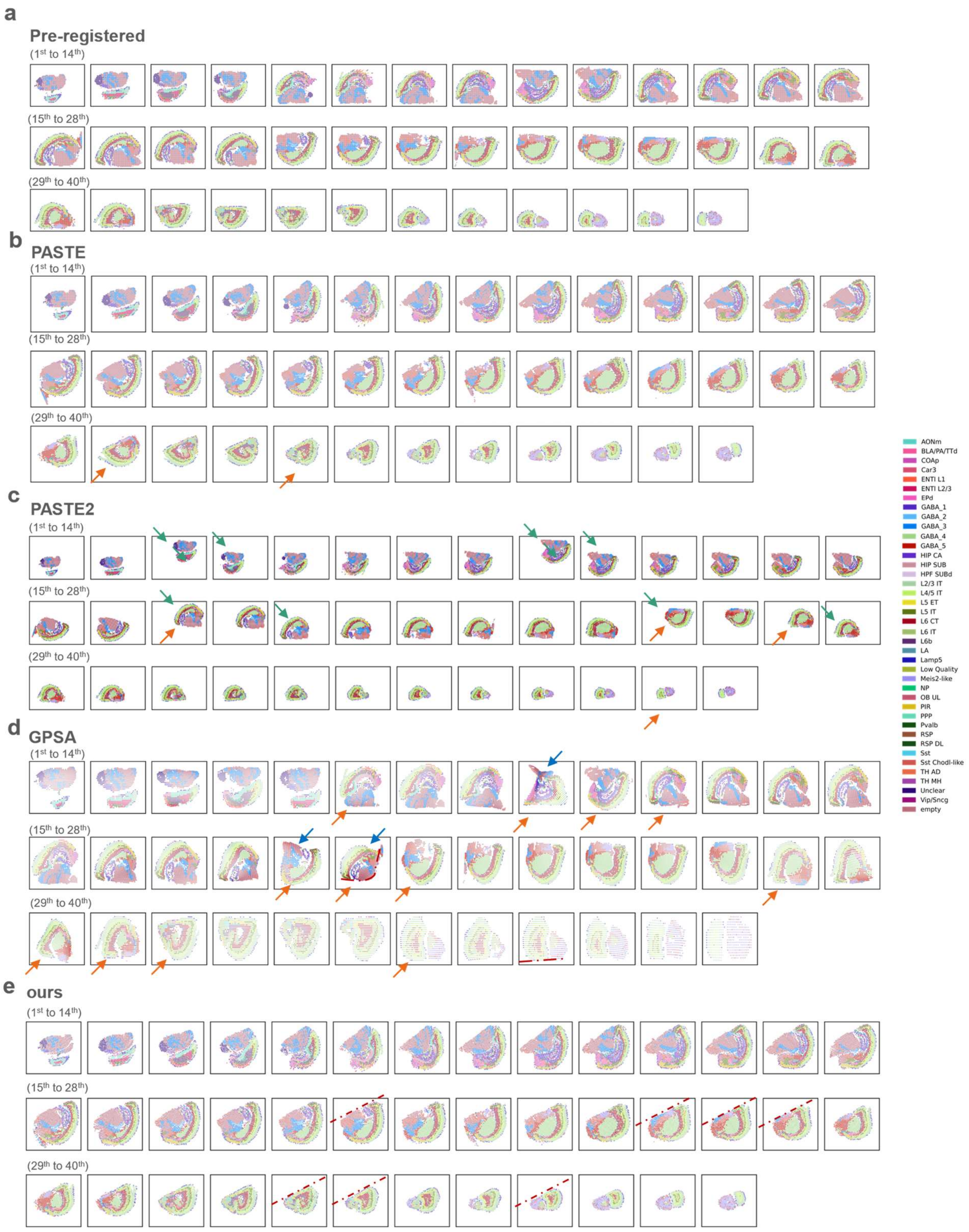


Supplementary Fig. 19: **ST-GEARS correctly registers all sections of Mouse brain hemisphere, in contrast to PASTE, PASTE2 and GPSA.** Shown here are all individual 40 sections of, (a) before and after registration by (b) PASTE, (c) PASTE2 (d) GPSA and (e) our method. Positional misalignments are marked by arrows of green, and angular misalignments are marked by arrows of orange. Visible cutting lines used to check angular alignment of result of our method, and mistaken shape distorsions by GPSA are marked by dotted lines. While positional misalignments are found in result of PASTE2, and angular misalignments are found in results both of PASTE2 and PASTE, neither of the conditions are visible in result of ST-GEARS.

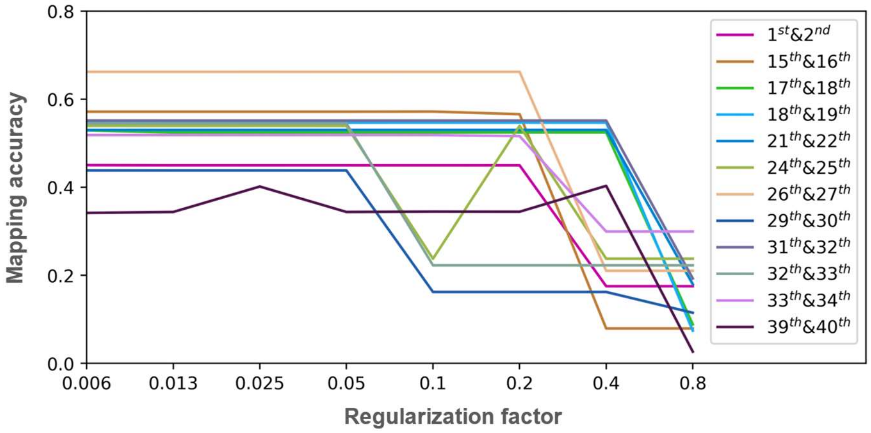


Supplementary Fig. 20: **Self-adaptive regularization generates regularization factor in Mouse brain registration that causes most advanced anchor accuracy.** The figure shows changing mapping accuracy of anchors upon exponentially changing regularization factor. Section pairs with mapping accuracy changes over range of 0.1 are selected and plotted. ST-GEARS runs through the regularization factors, and adopts the factor that causes highest mapping accuracy to ensure a most advanced anchor accuracy.
